# Diet outperforms microbial transplant to drive microbiome recovery post-antibiotics

**DOI:** 10.1101/2024.08.01.606245

**Authors:** M. Kennedy, A. Freiburger, M. Cooper, K. Beilsmith, M.L. St. George, M. Kalski, C. Cham, A. Guzzetta, S.C. Ng, F.K. Chan, D. Rubin, C.S. Henry, J. Bergelson, E.B. Chang

## Abstract

High-fat, low-fiber Western-style diets (WD) induce microbiome dysbiosis characterized by reduced taxonomic diversity and metabolic breadth^1,2^, which in turn increases risk for a wide array of metabolic^3–5^, immune^6^ and systemic pathologies. Recent work has established that WD can impair microbiome resilience to acute perturbations like antibiotic treatment^7,8^, although we know little about the mechanism of impairment and the specific host consequences of prolonged post-antibiotic dysbiosis. Here, we characterize the trajectory by which the gut microbiome recovers its taxonomic and functional profile after antibiotic treatment in mice on regular chow (RC) and WD, and find that only mice on RC undergo a rapid successional process of recovery. Metabolic modeling indicates that RC diet promotes the development of syntrophic cross-feeding interactions, while on WD, a dominant taxon monopolizes readily available resources without releasing syntrophic byproducts. Intervention experiments reveal that an appropriate dietary resource environment is both necessary and sufficient for rapid and robust microbiome recovery, whereas microbial transplant is neither. Furthermore, prolonged post-antibiotic dysbiosis in mice on WD renders them susceptible to infection by the intestinal pathogen *Salmonella enterica* serovar Typhimurium. Our data challenge widespread enthusiasm for fecal microbiota transplant (FMT) as a strategy to address dysbiosis and demonstrate that specific dietary interventions are, at minimum, an essential prerequisite for effective FMT, and may afford a safer, more natural, and less invasive alternative to FMT.

## INTRODUCTION

An extensive body of work shows that Western-style diets (WD) with high fat and refined sugar content and low dietary fiber promote gut dysbiosis characterized by reduced taxonomic diversity and altered function, with significant impacts on host metabolic, immune, and other organismal outcomes^1,2^. Recent data shows that diet-induced dysbiosis predisposes the microbiome to collapse after antibiotic perturbation^7,8^, but the relative contributions of diet and microbial community structure to this phenomenon are unknown, as is the extent to which the resultant dynamics can be explained by metabolic interactions. Microbial dysbiosis may reduce the availability of microbes to repopulate^7,9,10^, while a poor resource environment may alter the ecology of recovery and community diversification^11,12^. Understanding these dynamics is essential for choosing the most appropriate strategy for microbiome restoration. Here, we compare microbiome robustness to and recovery after antibiotic perturbation across host diets, and we evaluate interventions that allow us to disentangle the impacts of diet and microbial re- seeding on microbiome recovery.

## RESULTS

### Microbiome recovery across diets

To determine how diet impacts gut microbiome resilience to antibiotic treatment, 86 female specific-pathogen-free (SPF) C57Bl/6 mice underwent gut microbiota homogenization^13^ and were then acclimated to either a standard, low-fat, high-fiber diet (regular chow, “RC”) or a high-fat, low-fiber Western-style diet (“WD”) for 4 weeks. We then treated mice on each diet with either a triple antibiotic cocktail (“ABX”) or 5% PBS control in the drinking water for 72 hours and collected fecal samples through 4 weeks (Figure 1A, Materials and Methods). To assess long-term recovery, fecal samples were collected from 6 of these mice on both RC-ABX and WD-ABX through 9 weeks post-ABX. One additional cohort of 12 male mice split evenly across RC-ABX and WD-ABX underwent the same protocol through 2 weeks post-ABX to confirm that recovery patterns were consistent across sexes (Figure S1). Given that they were, we present analyses including only female cohorts unless otherwise specified.

**Figure 1:**
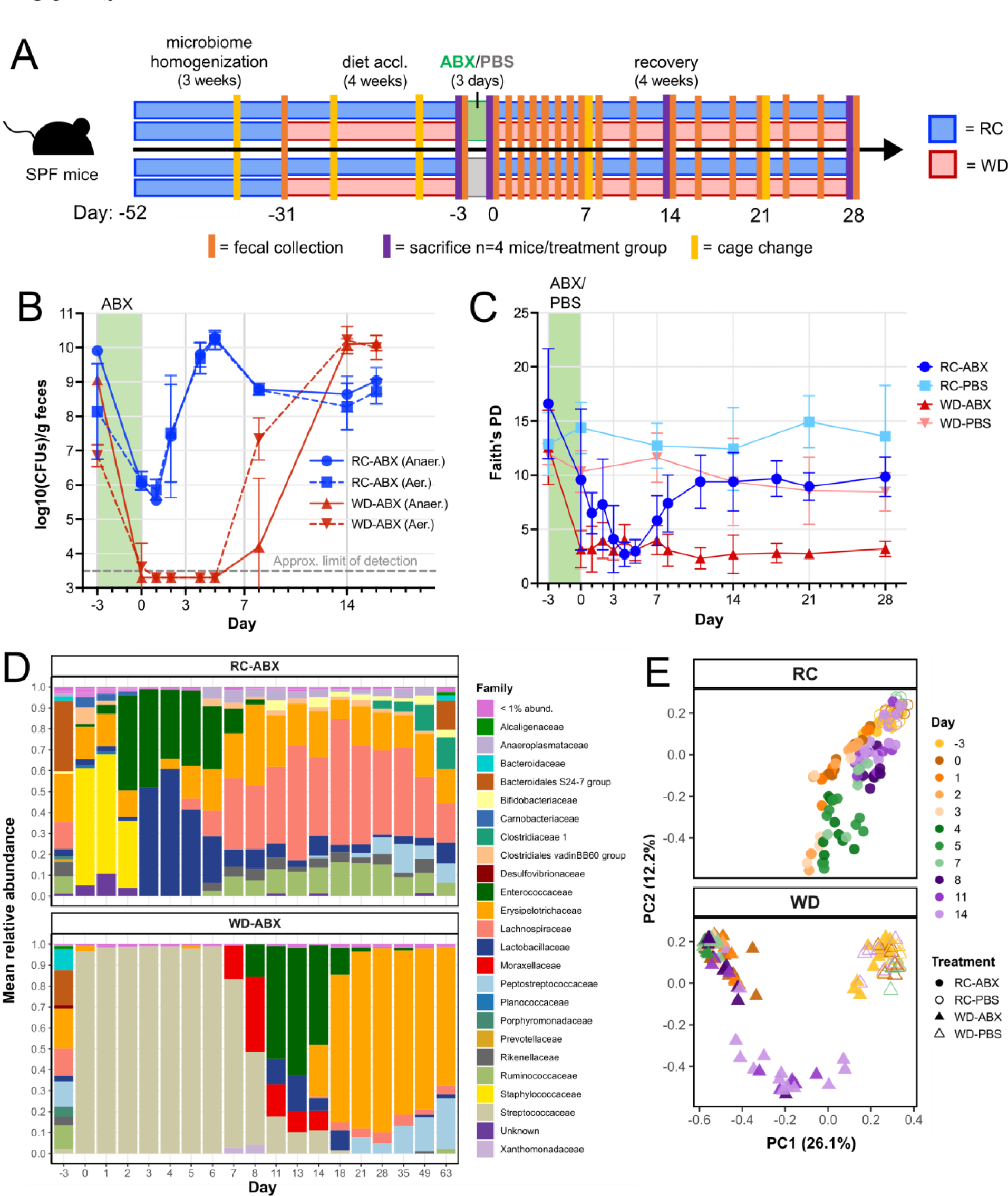
Bacterial biomass and taxonomic recovery after antibiotic treatment are impaired in mice on WD. (A) Mice on RC or WD were treated with PBS or ABX in the drinking water for 72 hours, and serial fecal samples were collected to assess microbiome recovery (Materials and Methods). (B) Fecal microbial biomass in mice on RC-ABX and WD- ABX for mice from Cohort 1 (n=6/group). See also Figure S1A. Error bars indicate mean ± SD. Statistics including exact *n* and *P* values are presented in Tables S1. (C) Fecal alpha diversity (phylogenetic diversity) of mice across dietary treatments and timepoints (all cohorts, n=4- 13/group). Error bars indicate mean ± SD. Statistics including exact *n* and *P* values are presented in Tables S2. (D) Mean relative abundances of different microbial families for Cohort 1 (n=6 mice/group, See Fig. S1F for other cohorts). (E) PCoA of 16S-based microbiome taxonomic composition at the genus level using Bray-Curtis dissimilarity for samples from all treatment groups and cohorts through Day 14. See Fig. S1G for results through Day 28.

We first quantified gut microbiome resilience to antibiotic treatment in terms of microbial biomass by counting colony-forming units (CFUs) cultured aerobically and anaerobically on rich media. We found that CFU counts dropped precipitously after antibiotic administration in both treatments, but much more severely for mice on WD (Figure 1B, Table S1). Moreover, while CFU counts for mice on RC had recovered to baseline levels by Day 4 post-ABX, CFU counts for mice on WD did not recover to baseline through at least Day 7. Microbial biomass of PBS controls remained stable in both dietary treatments (Figure S1B, Table S1).

We next evaluated taxonomic recovery of the gut microbiome across treatment groups using 16S rRNA gene sequencing. Immediately prior to antibiotic treatment (Day -3), mice on WD exhibited significantly reduced phylogenetic diversity compared to mice on RC (Figure 1C, Table S2). Antibiotic-treated mice on both diets experienced a sharp decline in phylogenetic diversity after antibiotic treatment. RC-ABX mice began to recover phylogenetic diversity after Day 5, and recovered over half the diversity that was initially present by Day 11. For WD-ABX mice, phylogenetic diversity remained severely diminished through at least Day 28 in all cohorts, and up to 9 weeks post-ABX. Phylogenetic diversity in PBS controls did not change significantly. Other metrics of alpha diversity (ASV richness, Shannon index) recapitulated these trends (Figure S1D-F, Table S2).

Correspondingly, relative abundances of taxa were substantially altered during and after antibiotic treatment for mice on both diets (Figure 1D). In RC-ABX mice, the microbiota passed through successive stages of recolonization after antibiotic treatment, marked by early dominance of facultative anaerobes like *Enterococcaceae* and *Lactobacillaceae*, followed by increasing diversification of stricter anaerobes. In WD-ABX mice, the low-biomass post-ABX community was dominated by *Streptococcaceae* until biomass started to recover. Recovery in WD-ABX mice also passed through a successional phase of facultative anaerobes including *Moraxellaceae*, *Enterococcaceae*, and *Lactobacillaceae* around Day 14, before recovering stricter anaerobes. While not all specific compositional changes were precisely replicated across cohorts (Figure S1G), certain characteristics, like the early dominance of facultative anaerobes as biomass recovers and the post-ABX dominance of *Lactococcus* within the *Streptococcaceae* family in all WD-ABX groups, were consistent.

PCoA of Bray-Curtis dissimilarity revealed that microbiome dynamics track a broadly similar trajectory within treatment groups across all cohorts (Figure 1E, Figure S1H-I). By Day 14 or earlier, the gut microbiota of mice on RC-ABX approached their respective pre-ABX community and corresponding RC-PBS controls, whereas the gut microbiota of WD-ABX mice remained distinct from their pre-ABX community at Day 14. By Day 28, while the microbiota of RC-ABX mice was indistinguishable from RC-PBS controls, only some mice on WD-ABX began to approach their initial community structure (Table S2). Together, these results indicate that mice on WD experience markedly impaired recovery of gut microbial taxonomic and biomass recovery after antibiotic treatment compared to mice on RC.

### Microbiome functional capacity

To determine whether the gut microbiota of mice with incomplete taxonomic recovery also exhibit altered functional capacity, we performed shotgun metagenomic sequencing on fecal samples from a subset of mice on RC-ABX and WD-ABX at key time points before antibiotic treatment and throughout recovery (Materials and Methods). Gene calls were annotated with the Kyoto Encyclopedia of Genes and Genomes (KEGG) catalog for functional interpretation.

We first evaluated gut microbiome functional diversity by calculating functional richness at different hierarchical levels (gene call, KEGG Ortholog (KO), or KEGG Category (KCat)) (Figure S2A). At the level of gene calls, mice on WD had reduced functional richness compared to mice on RC even before antibiotic treatment (Table S3). After antibiotics, functional richness of mice on both diets collapsed severely, and while mice on RC recovered up to 69% of their initial gene count by Day 28, mice on WD recovered only 16%. This mirrors the taxonomic trends observed in Figure 1.

At broader hierarchical levels like KO or KCat, functional richness was more preserved during and after antibiotic treatment across both treatment groups (Figure S2A). This could indicate a change in functional redundancy, i.e. the number of unique gene calls that map to each KO, before and after antibiotic treatment. High functional redundancy could permit robustness at the KO-level to the loss of individual gene calls. Indeed, we observe that mice on RC have greater functional redundancy before antibiotics and recover significantly more functional redundancy afterwards, whereas the loss of functional redundancy in mice on WD persists (Figure 2A, Table S3). In mice on RC, there was a much stronger correlation between initial and final functional redundancy across KOs than in mice on WD (Figure S2B, Table S3), which tended to lose functional redundancy irrespective of how much redundancy a given KO began with. To assess whether recovery of functional redundancy differed across functional subsystems, we mapped KOs with strong (>75%) or poor (<25%) recovery of functional redundancy at the KEGG system level (Figure 2B). Although there were many more KOs that exhibited poor recovery in mice on WD and the majority (55%) of these mapped to metabolic functions, the proportional breakdown of KEGG systems was nearly identical across treatment groups and recovery levels (Figure 2B). Thus, mice on WD experience a loss in functional diversity and redundancy upon antibiotic treatment that varies in intensity across KO groups, with major losses in metabolic redundancy.

**Figure 2:**
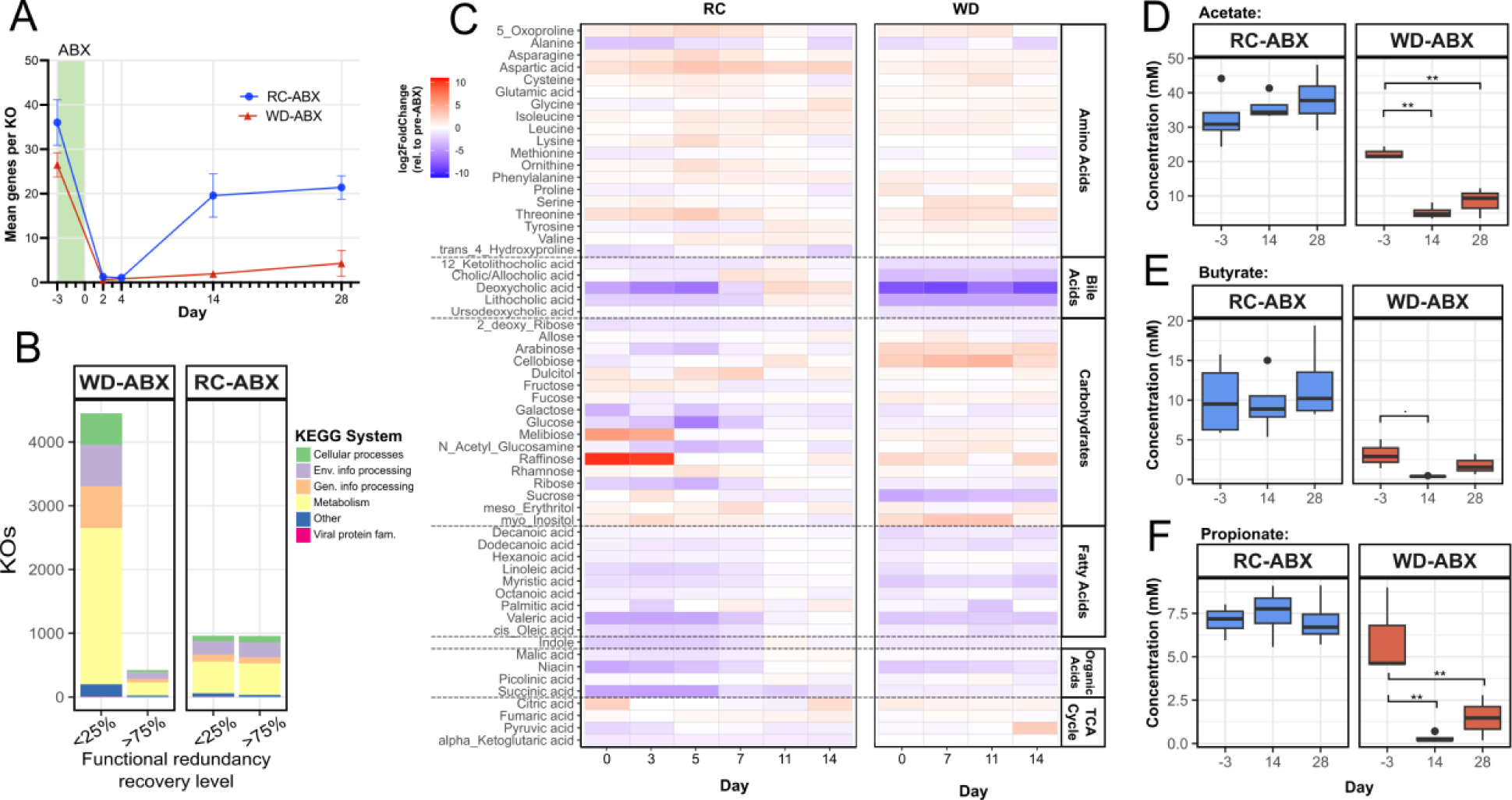
Functional recovery is severely impaired in mice on WD. (A) Functional redundancy (mean genes per KO) of mice on RC-ABX and WD-ABX (n=2-8/group). Error bars indicate mean ± SD. Statistics including exact *n* and *P* statistics are presented in Table S3. (B) KEGG system mapping of KOs that recovered < 25% or > 75% of their pre-ABX Day -3 functional redundancy across RC-ABX and WD-ABX groups. (C) Heatmap displaying log2FoldChange in metabolite abundances relative to the pre-ABX Day -3 timepoint, averaged across n=3-6 mice/group. (D – F) Absolute concentrations of (D) acetate, (E) butyrate, (F) propionate in mice on RC-ABX (blue) and WD-ABX (red) (n=3-4/group, *q < 0.05, **q < 0.01). Whiskers represent median ± 1.5*IQR. Statistics including exact *n* and *P* values are presented in Table S5.

In order to focus on the most functionally interpretable aspects of our data, all subsequent metagenomic analyses were performed at the KO level. We performed pairwise comparisons between the pre-ABX timepoint and all post-ABX timepoints to investigate functional characteristics of the microbiome throughout recovery, and to identify differentially abundant KOs. Mice on RC had 658 significantly depleted KOs at Day 2 relative to pre-ABX, which grew to 835 depleted KOs at Day 4, but nearly all of these recovered to baseline levels by Day 14 (Figure S2C). At Day 2 and Day 4, semi-overlapping but distinct subsets of genes were enriched or depleted (Figure S2D-E), indicating unique intermediate functional stages during recovery.

Mice on WD had a larger number of significantly depleted KOs than RC counterparts at all timepoints evaluated, and even by Day 28, 291 KOs had not yet returned to pre-ABX levels (Figure S2F). In contrast to mice on RC, the KOs depleted at Day 14 and Day 28 were almost all a subset of the KOs depleted at Day 2 (Figure S2G). Mice on WD also had small, semi-overlapping sets of KOs that were significantly enriched relative to pre-ABX at all timepoints (Figure S2H). Further analysis of specific functional representations across timepoints and diets are presented in Figure S2I-N and Table S4).

### WD impairs metabolome recovery

To directly assay the resource environment of the gut, we performed a targeted fecal metabolomic screen of gut microbiome-associated compounds including amino acids, carbohydrates, bile acids (BAs), and more, from mice on RC-ABX and WD-ABX before and after antibiotic treatment. As normalized metabolite abundances and dynamics were broadly consistent across individual mice within dietary treatments (Figure S3A), normalized abundance values were averaged across mice for visualization (Figure 2C).

We found that for mice on RC, the normalized abundances of many compounds were distinct from baseline immediately after antibiotics, but that by Day 11, the profile returned to baseline. This was statistically confirmed by PCoA clustering analysis (Figure S3B-C, Table S5). Our heatmap and PCoA revealed two inflection points during recovery: the first after Day 3 and the second after Day 7. The shift after Day 3 is driven largely by the dynamics of raffinose and melibiose, two plant-derived α-galactoside compounds that are highly abundant in the RC diet fed to our mice^14^. These compounds were highly elevated after antibiotics and through Day 3 but returned to baseline by Day 5. The second shift reflects the dynamics of many compounds – including carbohydrate monomers like ribose, glucose, and arabinose, as well as most fatty acids assayed - that are depleted after antibiotics but recover between Day 7 and Day 11. The secondary BAs lithocholic acid, deoxycholic acid, and 1,2-ketolithocholic acid, and to a lesser extent, the primary BA cholic acid were also depleted through Day 7, but were more abundant than baseline by Day 11. In stark contrast, the metabolomic profile of mice on WD showed almost no signs of recovery through Day 14 (Figure 2C, Figure S3B, C). Bile acids stood out as heavily depleted throughout sampling. Sucrose, niacin, and fatty acids, all of which were initially depleted in mice on RC but recovered after Day 7, remained depleted throughout sampling for mice on WD. Carbohydrates like cellobiose, arabinose, and myo-inositol were overly abundant relative to baseline and failed to return to baseline levels.

Because short-chain fatty acids (SCFAs) are well documented products of microbial metabolism derived from the breakdown of complex polysaccharides and can directly impact the host, we performed separate metabolomic analyses to quantify absolute levels of SCFAs in cecal samples from mice on RC-ABX and WD-ABX. The concentrations of acetate, butyrate, and propionate at Day 14 and Day 28 in mice on RC were statistically indistinguishable from pre-ABX, whereas for WD mice, they were depleted at Day 14 and persisted at low levels through Day 28 (Figures 2D-F, Table S5).

We wondered if changes in select metabolite abundances might correspond to changes in abundances of microbial genes known to produce or degrade those compounds. To assess this possibility, we plotted the relative abundances of several curated subsets of microbial genes across timepoints for mice on each dietary treatment and overlaid the abundances of the associated metabolites (Materials and Methods). For example, melibiose and raffinose contain an ⍺-1,6 linkage that can be broken by bacterial ⍺-galactosidase genes but not by host enzymes^14^. In mice on RC, melibiose and raffinose reach higher concentrations after antibiotics when microbial ⍺-galactosidase genes are most depleted, and as these genes recover, melibiose and raffinose abundances fall (Figure S3D, Table S6). In mice on WD, we observe few changes in the abundance of ⍺-galactosidase genes or the abundance of melibiose or raffinose. Starch and arabinan can similarly be metabolized into glucose and arabinose monomers, respectively. In mice on RC, both glucose and arabinose reach low abundance when genes for starch and arabinose metabolism drop, and as those polysaccharide metabolism genes rise in abundance, so does the abundance of the respective monomeric breakdown product (Figure S3E, F). In mice on WD, there are again few changes in either metabolite or gene abundances over the course of recovery. These data suggest that in mice on RC diet, the microbiota may respond to or interact with the resource environment, and especially with complex carbohydrates like those analyzed here, in a way that the microbiota of mice on WD does not.

### Metabolic modeling of recovery dynamics

Across all taxonomic and functional metrics evaluated, mice on WD experienced more severe ecosystem collapse with slower and less complete recovery than mice on RC (Figures 1- 2). This was not attributable to slower antibiotic clearance in mice on WD (Figure S4, Table S7). To explore the mechanism by which resource environment shapes our observed community recovery patterns, we developed metabolic models that leverage functional knowledge of microbiome members and integrated ‘omics data to predict the metabolic interactions and dynamics of each community over time (Materials and Methods). Most of the ASVs presented in Figure 1 closely match 16S sequences found in at least one fully annotated isolate genome in RefSeq, which we used to reconstruct representative probabilistic genome-scale metabolic models (prGEMs) through the ModelSEED2 pipeline in KBase. We simulated these ASV-based prGEMs to calculate the probability that each strain can take up, grow on, or excrete the compounds in our metabolomics assay, producing strain-metabolite interaction probability profiles (SMIPPs). Clustered heatmaps of these SMIPPs (Figure S5) suggest metabolic specialization among certain ASVs, and create a roadmap for understanding how each ASV can metabolically interact with others in the community.

We then combined the SMIPPs with our16S-based ASV-abundance profiles to compute microbiome-metabolite interaction probability profiles (MMIPPs) for each sample. These MMIPPs predict sample-wide metabolite consumption, conversion, and production capacity by component ASVs. Correlations of MMIPPs with measured metabolite abundances (Table S8) reveal that melibiose and cellobiose have higher correlations with predicted microbial consumption and production: these metabolites are abundant in samples predicted to produce the compounds, and scarce in samples predicted to consume them. This suggests interdependence between the dynamics of these compounds and microbiome metabolism.

To better understand which metabolic functions were performed by which ASVs over the course of recovery and how this differed across diets, we next combined our ASV-based prGEMs into community models and, for each interval between timepoints, simulated the flux through each metabolic pathway within the constraints of our observed metabolite and ASV abundance dynamics. These simulations determined the most likely metabolic behavior of the ASVs in each interval (Figure 3A, Table S8), and revealed profound metabolic differences between the RC and WD treatment groups. In the RC communities, ecological complexity and syntrophic cross-feeding interactions are maintained in the immediate aftermath of antibiotic treatment and throughout recovery (Figure 3B, Figure S6). By contrast, in WD communities, almost all syntrophic interactions are lost in the aftermath of antibiotic treatment through Day 7- 11. Instead, the WD communities are dominated by a single Lactococcus ASV with broad metabolic capacity that produces few syntrophic byproducts. The RC community contains this same Lactococcus ASV, but in the RC dietary context, it is balanced by trophic complexity and interdependencies with other strains, and never dominates the microbiome. Although amino acids are excluded from Figure 3A for visual clarity, the observed patterns of greater metabolic interactivity, syntrophy, and complexity in the RC microbiome compared to WD apply to amino acid metabolism as well (Figure 3B, Table S8).

**Figure 3:**
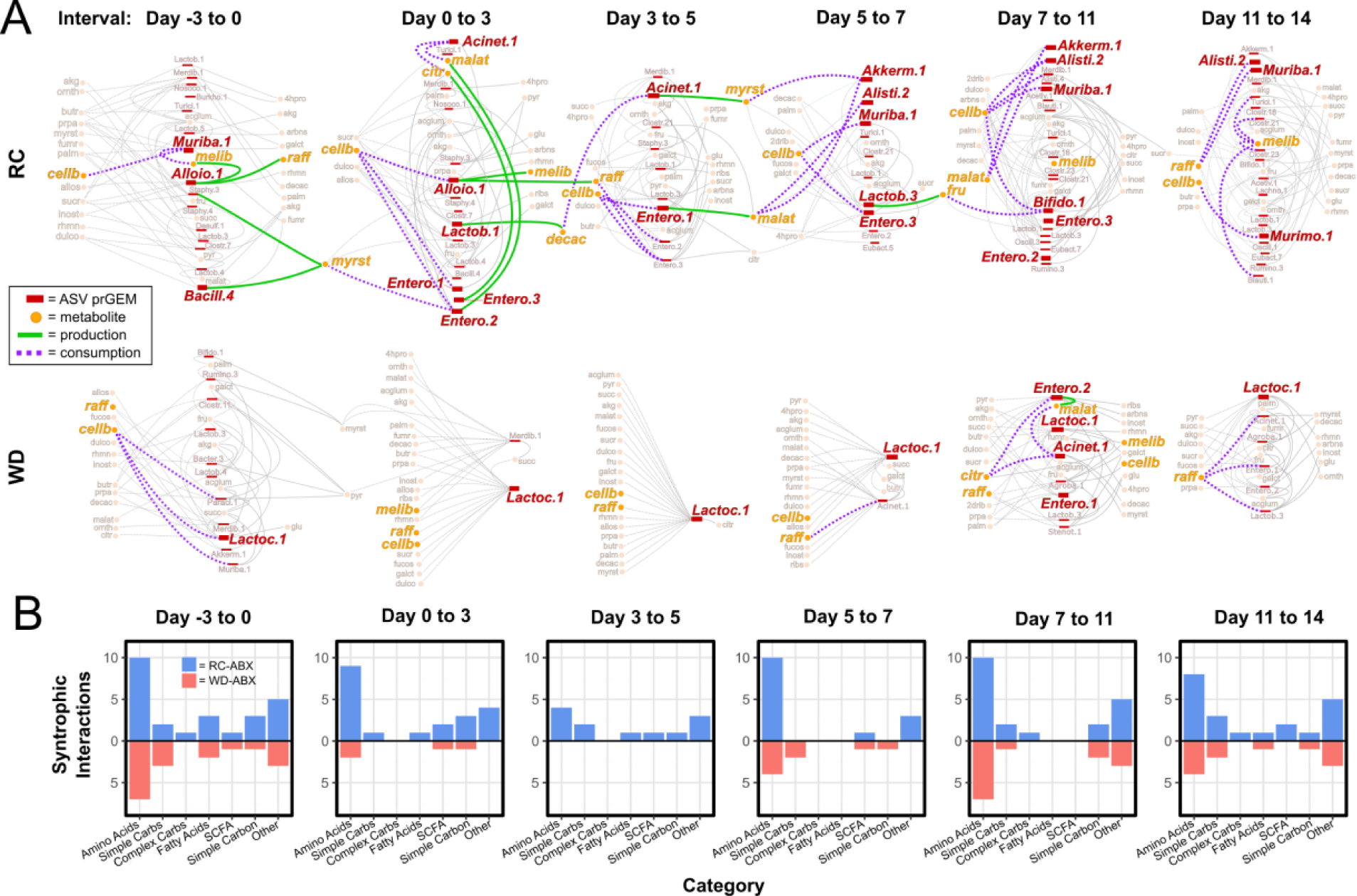
**Metabolic modeling predicts poor syntrophy in mice on WD**. (A) Community flux simulations over each time interval for mice on RC-ABX or WD-ABX. Edges represent predicted flux (dashed = consumption, solid = production) and nodes represent metabolites (orange) or ASV prGEMs (red). Select fluxes predicted to play a crucial role in recovery are highlighted in green and purple; other fluxes are grey. Map includes all measured metabolites with fluxes > 0.05 excluding niacin, acetate, propionate, and amino acids for visual clarity, but full model with all flux values is available in Table S8. Interactive map with flux values is available at https://modelseed.org/annotation/projects/gut_microbiome/. (B) Number of syntrophic interactions identified in the community models for mice on RC (blue) and WD (red) by category over each time interval.

The community simulations further reveal a pattern of metabolic succession in the recovery of the RC microbiome. Muribaculum abundance declines following antibiotic treatment, leaving open a niche for the consumption of cellobiose. This is filled by various Enterococci, which consume cellobiose and raffinose and produce metabolic products like citrate and malate, and thereby cultivate a niche for Acinetobacter to emerge. The continued production of malate from Enterococcus and the production of myristate from Acinetobacter open a niche for Akkermansia, and the complex carbohydrate consumers Alistipes and Muribaculum. By Day 7-11, Muribaculum has largely replaced Enterococcus as the primary consumer of cellobiose and raffinose, as in the pre-antibiotic state. The partial recovery observed in WD mice between Day 7 -14 also seems to be mediated by the emergence of Enterococcus. In this dietary context however, they are insufficient to shift the community towards complete recovery.

Collectively, our model suggests that metabolism of complex carbohydrates like cellobiose and raffinose, which are relatively more abundant in RC than in the purified WD, drive the syntrophic interactions that facilitate succession, diversification, and recovery. On WD, despite the microbiome’s overlapping taxa and relatively broad capacity to metabolize the same compounds after antibiotics, the greater availability of simple sugars promotes dominance of a single taxon. Thus, dietary resource availability fundamentally shapes the way that available taxa interact with their environment and other microbes to promote or prevent recovery.

### Dietary intervention versus FMT

Our data suggest that recovery in WD is limited primarily by an imbalance in the availability of simple and complex carbohydrates across dietary treatments rather than lack of metabolically capable taxa. To test this, we performed intervention experiments in which diet and microbial re-exposures were controlled after antibiotic treatment (Figure 4A, Materials and Methods). Briefly, after antibiotic treatment, mice on each pre-antibiotic diet were transferred into sterile gnotobiotic cages, and different post-antibiotic diet (“RC_D_”, “WD_D_”) and/or microbial re-exposures (“RC_M_”, “WD_M_”) were administered in a factorial manner: for mice on each pre-antibiotic diet, we changed either microbial re-exposure, diet, both, or neither. Microbial re-exposure was administered via FMT at 24 hours post-antibiotics and again at Day 14; one group on each diet received sterile PBS_M_ gavage as a no-transplant control. Fecal samples were collected before and after antibiotic treatment, and weekly through Day 28 of recovery. We selected Day 14 as the most salient timepoint to evaluate the distinct recovery dynamics of mice on RC and WD (full time course data available in Figure S7, Supplementary Discussion). No-ABX controls on each diet serve as benchmarks for “recovery,” which was evaluated in terms of overall community composition via 16S rRNA sequencing (Figure 4B) and further quantified by PCoA clustering analysis (Figure S7, Table S9) and ASV richness (Figure 4C).

**Figure 4:**
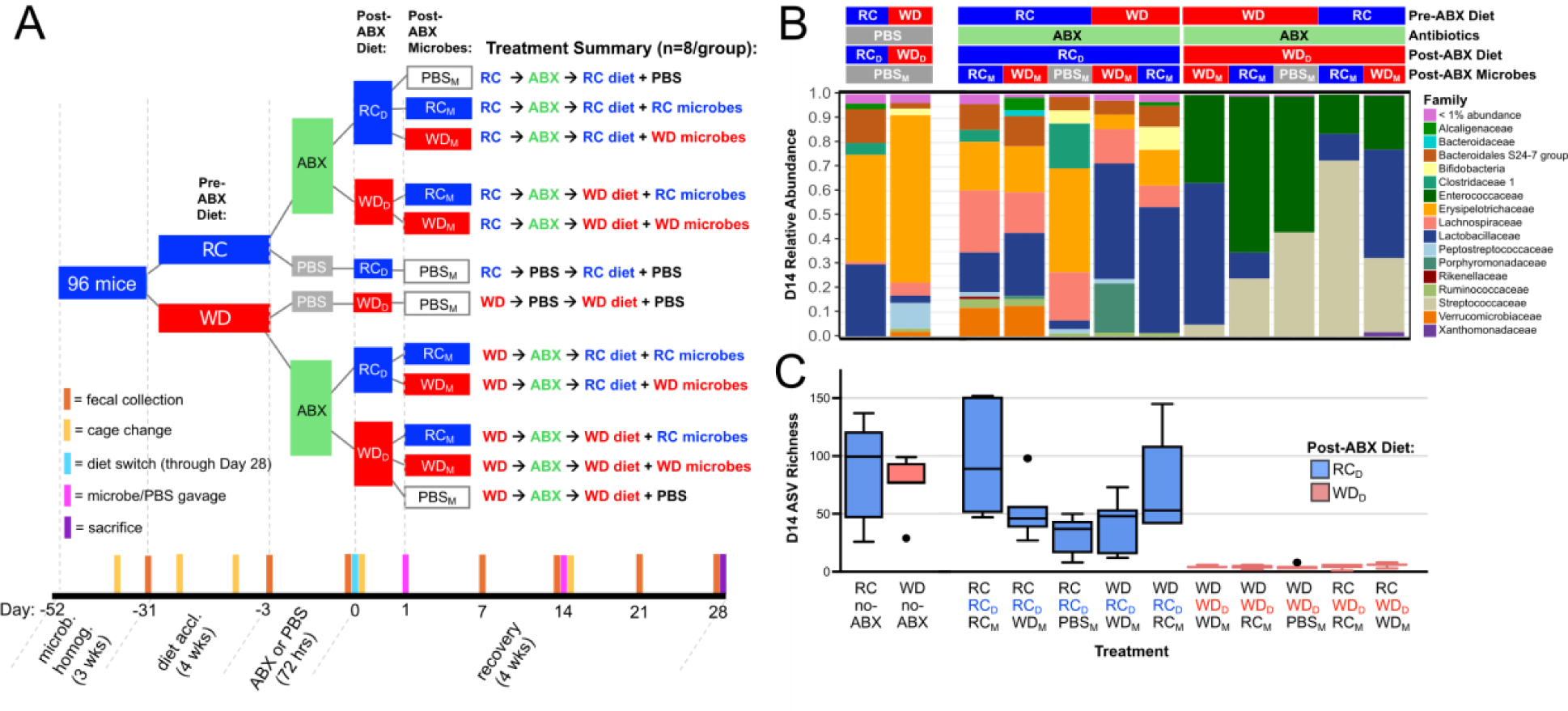
Dietary intervention facilitates microbiome recovery from antibiotics. (A) Experimental design. After ABX or PBS, post-ABX dietary treatments (RC_D_, WD_D_) were provided ad libitum from day 0 through the end of the experiment; post-ABX microbial treatments (RC_M_, WD_M_, PBS_M_) were administered at Day 1 and Day 14. (B) Mean relative abundances of microbial families at Day 14 across treatment groups (n=5-8 mice/group). (C) ASV richness across treatment groups at Day 14 (n=5-8/group). Whiskers indicate median ± 1.5*IQR. Statistics including exact *n* and *P* values are presented in Table S9.

We first confirmed that this experimental model recapitulates the phenotypes of our original experiments: mice that did not change dietary or microbial re-exposures (RC → RC_D_/RC_M_ and WD → WD_D_/WD_M_) matched the RC-ABX and WD-ABX phenotypes described in Figure 1 (i.e. RC recovered gut microbiota composition and alpha diversity, WD did not).

From relative abundance plots, we see that irrespective of pre-ABX diet or post-ABX microbial transplant, gut microbiota composition broadly segregates on the basis of post-ABX diet (Figure 4B). PCoA analysis statistically confirms that gut microbial composition segregates across PC1 by post-ABX diet, with mice on RC_D_ uniformly falling lower on PC1 and closer to the no-ABX controls than mice on WD_D_ (Figure S7, Table S9). Similarly, mice on post-ABX RC_D_ uniformly recovered more ASV richness by D14 than all WD_D_ counterparts (Figure 4C, Table S9).

Among mice that were fed post-ABX WD_D_, microbial transplant had negligible impact on recovery, with all WD_D_ treatment groups exhibiting severely diminished ASV richness and clustering distinctly from WD no-ABX controls at D14. These experiments indicate that an appropriate diet is both necessary and sufficient for rapid and robust gut microbiota recovery after antibiotic treatment, while microbial transplant is neither. This broadly supports our modeling predictions that recovery is predominantly driven by dietary resource availability (especially the balance of simple and complex carbohydrates) rather than by the presence or absence of specific taxa.

### Prolonged loss of colonization resistance

Under healthy conditions, the microbiome protects the host via “colonization resistance” against opportunistic pathogens^15^. For example, the pathogen *Salmonella enterica* serovar *Typhimurium* (*St*) is unable to establish lower GI infection or cause colitis in SPF mice unless they have been pre-treated with streptomycin^16^. While established models of *St* colonization resistance require pre-treatment with antibiotics for 24 hours immediately before *St* challenge, we wondered if the prolonged post-antibiotic dysbiosis experienced by mice on WD might render them more susceptible to opportunistic infection by *St* as late as 14 days after antibiotic treatment has ended.

To evaluate this possibility, we performed a series of experiments including 5 cohorts of female mice and one cohort of male mice in standard, non-gnotobiotic cages (Materials and Methods). After antibiotic or PBS administration, mice recovered for 14 days, as in other experiments, and then each dietary and antibiotic treatment group was split into infection and no-infection treatment groups (Figure 5A). WD has been shown to facilitate *St* infection relative to RC even without antibiotic pre-treatment^17^; by comparing ABX and no-ABX control groups on each diet, we can isolate the specific effects of diet versus diet-induced post-antibiotic dysbiosis. Fecal samples and body weights were collected at t=6, 12, 24, 48, 72, and 96 hours post-infection (hpi). As trends were consistent across male and female cohorts (Figure S8), data were combined to improve statistical power.

**Figure 5:**
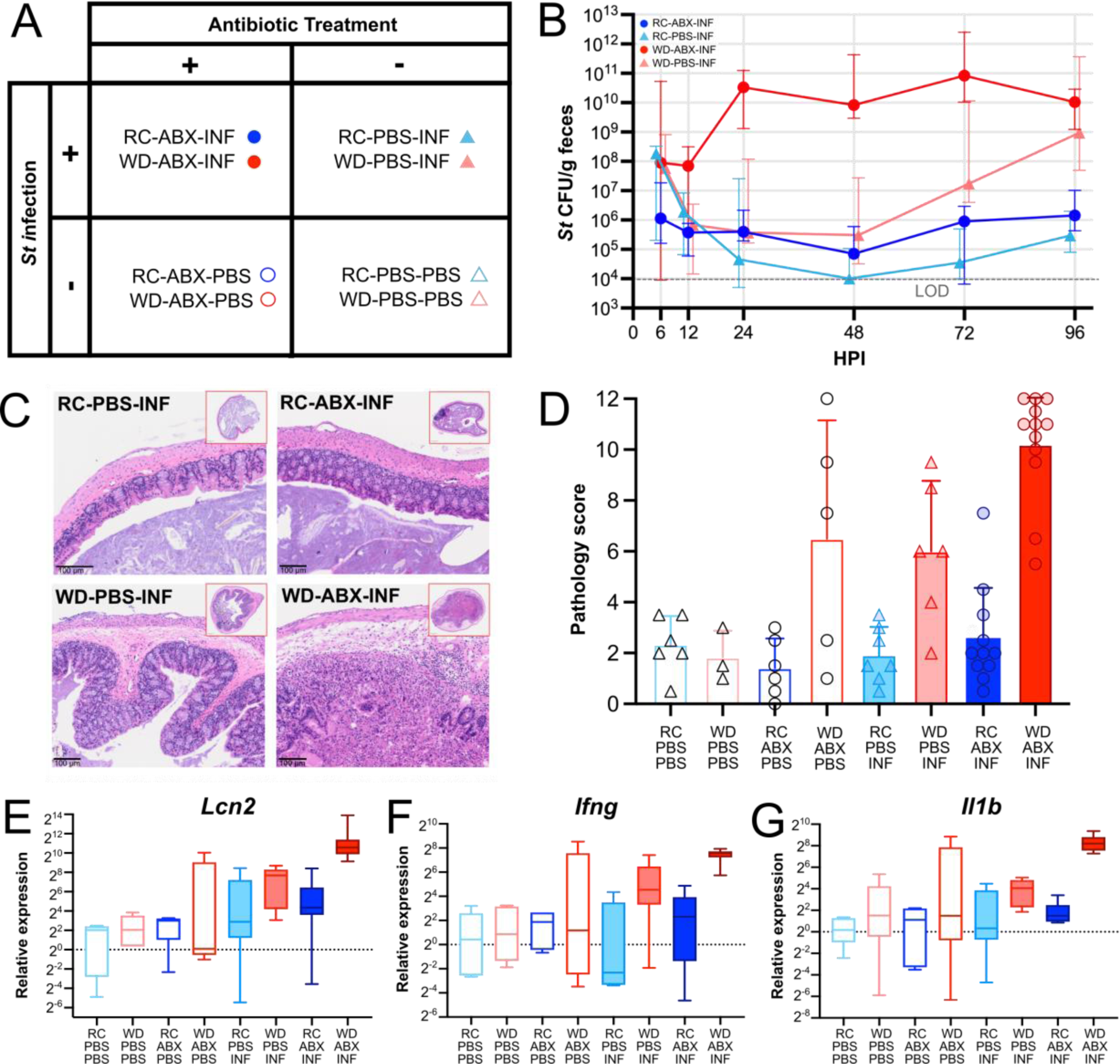
Prolonged post-antibiotic dysbiosis in mice on WD impairs colonization resistance to St. (A) Experimental treatment groups. (B) Fecal St load (CFU/g) among infected treatment groups. Uninfected controls had no detectable *St* and are not depicted. Error bars indicate median ± 1.5*IQR. Statistics including exact *n* and *P* values are presented in Table S10. (C) Representative histological images of mice in different treatment groups. (D) Histopathological scoring of cecal sections from mice on indicated treatment groups at 96 hpi (n=5–13/group, Figure S8). Error bars indicate mean ± SD. Statistics including exact *n* and *P* values are presented in Table S10. (E – G) mRNA expression of immune genes in cecal mucosal scrapings at t=96 hpi based on RT-qPCR (see Figure S8 for additional inflammatory markers). Expression is normalized to the housekeeping gene Actb and the RC-PBS-PBS treatment group (n=3-13/group). Whiskers indicate median ± 1.5*IQR. Statistics including exact *n* and *P* values are presented in Table S10.

We first evaluated *St* load at each timepoint after infection (Figure 5B). Uninfected controls had undetectable levels of *St*, confirming no contamination. As hypothesized, mice on WD-ABX-INF were the most susceptible to infection, exhibiting significantly higher *St* loads than all other treatment groups from t=24-48 hpi and reaching ∼10^5^-fold higher median infection load than the WD-PBS-INF group during this period (Table S10). From 72-96 hpi, both the WD- ABX-INF and WD-PBS-INF groups had significantly greater infection load than all groups on RC diet, recapitulating previous findings that WD alone is sufficient to impair colonization resistance. Although the WD-ABX-INF group had greater median *St* load than WD-PBS-INF through 96 hpi, this difference was no longer significantly different after 48 hpi.

We performed targeted analyses of infection severity in lower GI tissues to focus on the enteric colonization resistance (rather than disseminated typhoid) model of *St* infection (gross body weight data are available in Figure S8, Supplementary Discussion)^16,18^. Histopathological scoring and qPCR analysis of a panel of *St-*induced inflammatory markers in cecal tissue revealed that the WD-ABX-INF group experienced significantly more severe inflammatory pathology (mean histopathology score = 10.21 ± 2.126, Figures 5C, D, Figure S8, Table S10) with higher expression of inflammatory markers (Figures 5E-G, Figure S8, Table S10) than all other treatment groups. Interestingly, the uninfected WD-ABX-PBS control group experienced variable but occasionally severe inflammation (mean histopathology score = 6.5 ± 4.65) comparable to the WD-ABX-INF group, suggesting that antibiotic treatment in mice on WD can induce inflammation even without *St* challenge. Together, these results indicate that only the WD-ABX-INF group consistently succumbed to lower-GI infection and enteritis, supporting the hypothesis that prolonged post-antibiotic dysbiosis in mice on WD impairs colonization resistance in the lower GI tract relative to mice that were not pre-treated with antibiotics, or that were on RC diet.

## DISCUSSION

Our experiments reveal that WD and antibiotic treatment collectively cause more severe and prolonged microbial ecosystem collapse than observed for either factor alone. This collapse entails extreme loss of taxonomic diversity, as previously reported^7,8^, as well as reduced functional redundancy and a skewed metabolome. Metabolic modeling indicates that recovery in mice on RC is driven by the development of syntrophy as complex carbohydrates are metabolized and simpler byproducts open niche space in the community. Although the microbiota of mice on WD retains the metagenomic capacity to perform these same reactions after antibiotic treatment, the easy availability of simple dietary sugars promotes dominance by a single metabolic generalist and prevents progression along the same successional trajectory.

These findings are supported by our intervention experiments, in which we showed that without an appropriate dietary resource environment, microbial communities will fail to recover, irrespective of microbial transplant. Last, we show that prolonged post-antibiotic dysbiosis in mice on WD can extend the window of susceptibility to opportunistic pathogens like *St*. Given that our WD formulation is designed to reflect typical American consumption patterns^19^ and that antibiotics are liberally prescribed for an enormous array of pathologies^20,21^, these findings bear great clinical relevance. In humans, variability in microbiome recovery after antibiotics is well documented, but poorly understood^22–26^. Our data suggest that diet may play a central role in driving inter-individual differences in microbiome recovery and should be explicitly assessed in human studies. Moreover, we must consider the risks of prolonged and severe post-antibiotic dysbiosis for populations vulnerable to opportunistic infection, like immunocompromised individuals, or those experiencing chronic disease. For instance, severe cases of ulcerative colitis are often treated with antibiotics^27^. Individuals with ulcerative colitis also often maintain restrictive diets to prevent flares^28^. The combination of limited diet and antibiotic treatment may paradoxically predispose these individuals to exacerbated microbiome dysbiosis, which may feed back into the cycle of disease. Appropriate peri-antibiotic dietary interventions should be investigated across clinical contexts as a safe and affordable route to promote microbiome recovery when antibiotics must be used.

To date, the growing field of microbiome therapeutics has centered largely around microbial replacement strategies like fecal microbiota transplant (FMT)^29^, probiotics^30^, or live biotherapeutics^31^, which have shown widely variable efficacy across individuals and diseases. Our intervention experiments indicate that diet is more foundational to recovery than microbial re-seeding, and that without an appropriate dietary resource environment, microbial transplant is insufficient to promote recovery. Even under controlled lab conditions using autologous FMT of a mature, diverse community in genetically matched, diet-controlled recipients, the outcomes of microbial transplant were variable and ultimately dependent upon post-antibiotic diet. In contexts with more significant microbial extinction than in our experiments, microbial replacement may yet play an important role in recovery, as has been extensively shown^7,9,10^. However, our data suggest that unless the microbial transplant encounters an environment with the right resources to support engraftment, growth, and diversification in the recipient, its efficacy will be limited at best. Thus, concomitant dietary interventions may improve the consistency and efficacy of existing microbial transplant strategies.

Finally, we propose ecological succession as a novel paradigm for approaching microbiome restoration. Under this model, both the taxa present and their surrounding resource environment interact in an iterative feedback process to guide the progression of the community from one stage to the next^32,33^. Early arriving microbes may facilitate or inhibit the growth of later arriving microbes by producing metabolic byproducts for cross-feeding^34^ or by changing the resource environment (e.g. oxygen levels^35^, pH^36^, bile acid pool^37^). The community cannot proceed to the next stage if it is either missing the right taxa, or if the taxa do not have access to the right resources. Our data and simulations indicate that WD does not provide the right balance of simple and complex carbohydrates to initiate this successional process even when the right taxa are present. In this sense, FMT in mice on WD is akin to transplanting a mature forest into barren soil after a fire: the soil is unable to accommodate its growth. By using approaches like the metabolic model presented here, we can resolve the transitional dynamics between healthy and disease states and learn to support succession at each stage by matching specific dietary or microbial interventions to the present needs of the community. In this way, we can promote the environmental change necessary for the ecosystem to once again accommodate the growth of the climax community.

## Supporting information

Supplemental Information Titles

Supplementary Discussion

## MATERIALS AND METHODS

### Mice

C57Bl/6 mice at 5 weeks of age were purchased from The Jackson Laboratory barrier facility EM04 and co-housed in cages of 2-4 mice with pine shavings bedding in standard barrier facilities unless otherwise specified. All mice underwent a two-step microbiome homogenization protocol: 1) bedding was mixed across all cages twice a week from age 5-8 weeks leading up to the beginning of each experiment to reduce intra-cohort cage effects, and 2) mice were gavaged once at 6-weeks with fecal material banked from our SPF colony to reduce inter-cohort differences in microbiome composition^13^. Mice were fed autoclaved standard RC diet (LabDiets 5K67) during microbiome homogenization. Most experiments used only female mice to minimize variability introduced by sex, but one male cohort was included to ensure that phenotypes were consistent across sexes. This cohort was formally part of the colonization resistance experiment, which underwent an identical protocol to all other experiments through Day 14 of recovery. All mouse experiments were conducted in accordance with the University of Chicago Institutional Biosafety Committee and Institutional Animal Care and Use Committee.

### Diet Acclimation and Antibiotic Treatment

Mice at 8 weeks of age were either maintained on RC diet (LabDiets 5K67) or switched to WD (Envigo TD.97222)^38^ to acclimate for 4 weeks. This diet was designed to broadly reflect the nutritional intake of a Western population based on the Center for Disease Control and Prevention’s National Health and Nutrition Examination Survey^19^. During diet acclimation, bedding was mixed across cages within diet treatments twice a week. At 12 weeks of age, mice on each diet were treated with a sterile-filtered triple antibiotic cocktail of vancomycin (0.5 mg/ml), neomycin (1.0 mg/ml), and cefoperazone (0.5mg/ml) or 5% sterile PBS control in the drinking water for 72 hours. Water consumption was monitored during this time to ensure adequate treatment (Figure S1A).

### Western Diet Microbiome Resilience Experiments

Three female cohorts of mice were used in these experiments (including the long-term cohort described below). After treatment with antibiotics or PBS control, mice were maintained on their respective pre-antibiotic diets, and fecal samples were collected for 4 weeks after cessation of antibiotics on days -31, -3, 0, 1-8, 11, 14, 16, 18, 21, 23, 25, and 28. Cages were changed on days-38, -24, -10, 7, and 21. Subsets of mice were sacrificed at the pre-antibiotic (Day -3), post-antibiotic (Day 0), week two of recovery (Day 14), or week four of recovery (Day 28) timepoints (total n=4/timepoint/treatment group).

### Western Diet Microbiome Resilience – Long-term Recovery Cohort

One cohort of 8-week old female C57Bl/6 mice (n=6/group, RC-ABX and WD-ABX only) was bred in our animal facility for this experiment. Mice were co-housed in cages of 3 mice with pine shavings bedding in standard barrier facilities and did not undergo the microbiome homogenization protocol. These mice were acclimated to their respective dietary treatment for only 10 days. Although 10-days of diet acclimation was predicted to be sufficient for microbiome stabilization, which can occur within 4 days^39^, we lengthened the diet acclimation phase in subsequent cohorts to ensure that host physiological differences across dietary groups had stabilized^40^. During diet acclimation, bedding was mixed across cages within diet treatments twice a week. Antibiotic treatment was administered as described above. Fecal samples were collected for 9 weeks after cessation of antibiotics on days -31, -3, 0, 0.5, 1, 1.5, 2-8, 11, 14, 16, 18, 21, 28, 35, 49, and 63, and then all mice were sacrificed. Cage changes were performed on days -10, 7, 21, 35, and 49. Experimental duration was decreased after this cohort as microbiome recovery appeared to stabilize by Day 28 (Fig. S1).

### Post-antibiotic Intervention Experiments

Four cohorts of female mice were used in these experiments (n=8/treatment group, n=96 total). For each cohort, 24 mice at 5 weeks of age were split across 4 cages (n=6/cage) in standard barrier facilities for 3 weeks of microbiome homogenization. After microbiome homogenization, mice were transferred into 8 hermetically sealed gnotobiotic Techniplast IsoCage P Bioexclusion cages for diet acclimation and antibiotic treatment as described above (n=3/cage). Immediately following antibiotic treatment (Day 0), mice were transferred into 12 new sterile gnotobiotic cages and post-antibiotic diet and microbial re-exposure treatments were administered (n=2/treatment group/cohort). For microbial re-exposure treatments, fecal material was collected from all mice on the day before antibiotic treatment (Day -1), pooled by dietary treatment, resuspended at 60 mg/ml in 25% glycerol solution, and frozen at -80°C until administration.

Microbial re-exposures were administered by oral gavage of 200 µl of the respective fecal solution or sterile PBS at 24 hours after cessation of antibiotics (Day 1) and were re-administered at Day 14 of recovery after cage changes were performed to ensure that dispersal limitation did not impair microbiome recovery. Fecal samples were collected on days -31, -3, 0, 7, 14, 21, and 28, and mice were sacrificed at Day 28 post-antibiotics.

All procedures on the IsoCage P rack system from the time of antibiotic administration onward were performed using a modified sterile technique necessitated by the logistics of handling such a large number of treatment groups: one researcher donned sterile garb including two pairs of sterile gloves, the workspace was covered with a sterile drape, and another team member assisted in manipulation of the outside of the cages. The sterile team member performed all mouse manipulations without making contact with anything outside of the cage or sterile field. In between treatment groups, the sterile team member donned new sterile gloves, but otherwise continued to use the same garb.

### Colonization Resistance Experiments

After diet acclimation and antibiotic or PBS administration, 4 cohorts of female mice (n=3- 6/treatment group) and one cohort of male mice (n=6/treatment group, RC-ABX-INF and WD-ABX-INF only) were allowed to recover for 14 days, and were then inoculated by oral gavage with 200 µl of either nalidixic acid-resistant *Salmonella enterica* serovar Typhimurium (*St*, strain IR715)^41^ or PBS control. To prepare the gavage solution, *St* was grown aerobically in Luria Broth (LB) media at 30°C with shaking at 250RPM for 14 hours. Cultures were pelleted by centrifuging for 5 minutes at 4000*xg* and resuspended at 1:100 dilution in PBS (final infection dose: ∼7 x 10^7^ CFU/mouse). Fecal samples and body weights were collected for 4 days post-infection, and then mice were sacrificed.

### Tissue Harvest and Sample Processing

All mice were euthanized by CO_2_ asphyxiation and death was confirmed via cervical dislocation. After sacrifice, blood was collected by cardiac puncture and serum was isolated and stored at - 80°C. Liver, spleen, mesenteric lymph nodes, and mesenteric, gonadal, inguinal, and retroperitoneal fat deposits were weighed and split across samples that were preserved for histology, snap-frozen for RNAseq, and homogenized for CFU counts. The GI tract was dissected out, the cecum was weighed, and colon length was measured. Sections of the ileum, cecum, and colon were preserved for histology. Luminal contents from each of these sections were homogenized for CFU counting and snap-frozen for metabolomics/DNA extraction, and mucosal scrapings from each section were snap-frozen for RNAseq and/or RT-qPCR.

### CFU Counts

Pre-weighed fecal samples were suspended in 500ml of 25% glycerol solution, homogenized for 1 minute in a Mini-BeadBeater-96 (no beads, 2400 RPM), and serially diluted in PBS. Overall bacterial load was quantified by plating on Brain Heart Infusion-Supplemented (BHI-S) agar and incubating aerobically and anaerobically at 37°C for 24 hours. Measurements of overall bacterial load were collected for the pilot cohort, as well as all Colonization Resistance experiments.

Although we do not have bacterial biomass measurements for the two non-pilot cohorts of the Western Diet Resilience experiment, the Colonization Resistance experiments were carried out identically to these experiments through Day 14 post-ABX, and data from these timepoints may therefore be interpreted in the same manner. These data, collected from 5 separate cohorts and 48 mice, recapitulate the bacterial biomass dynamics of the RC-ABX and WD-ABX treatment groups reported in Figure 1B (Fig. S1B, C). Moreover, they indicate no significant loss of bacterial biomass over the course of the experiment in RC-PBS or WD-PBS controls.

To evaluate *St* load, pre-weighed fecal or tissue samples were homogenized in 25% glycerol solution as described above, and quantified by plating on LB agar with 25ug/ml nalidixic acid and incubating at room temperature for 24 hours.

### DNA Extraction

DNA was extracted using the QIAamp PowerFecal Pro DNA kit (Qiagen). Prior to extraction, samples were subjected to mechanical disruption using a bead beating method. Briefly, samples were suspended in a bead tube (Qiagen) along with lysis buffer and loaded on a bead mill homogenizer (Fisherbrand). Samples were then centrifuged, and supernatant was resuspended in a reagent that effectively removed inhibitors. DNA was then purified routinely using a spin column filter membrane and quantified using Qubit.

### 16S rRNA Sequencing

The V4-V5 region within the 16S rRNA gene was amplified using universal bacterial primers – 563F (5’-nnnnnnnn-NNNNNNNNNNNN-AYTGGGYDTAAA- GNG-3’) and 926R (5’-nnnnnnnn-NNNNNNNNNNNN-CCGTCAATTYHT- TTRAGT-3’), where ‘N’ represents the barcodes, ‘n’ are additional nucleotides added to offset primer sequencing. Approximately∼412bp region amplicons were then purified using a spin column-based method (Qiagen), quantified, and pooled at equimolar concentrations. Illumina sequencing-compatible Unique Dual Index (UDI) adapters were ligated onto the pools using the QIAseq 1-step amplicon library kit (Qiagen). Library QC was performed using Qubit and Tapestation and sequenced on Illumina MiSeq platform to generate 2x250bp reads.

### Shotgun Metagenomics

Libraries were prepared using 100 ng of genomic DNA using the QIAseq FX DNA library kit (Qiagen). Briefly, DNA was fragmented enzymatically into smaller fragments and desired insert size was achieved by adjusting fragmentation conditions. Fragmented DNA was end repaired and ‘A’s’ were added to the 3’ends to stage inserts for ligation. During ligation step, Illumina-compatible UDI adapters were added to the inserts and the prepared library was PCR amplified. Amplified libraries were cleaned up, and QC was performed using a tapestation. Libraries were sequenced on an Illumina NextSeq 500 to generate 1x150 reads.

### Metagenomic Analysis

Raw metagenomics reads were trimmed using Trimmomatic^42^, and a Minoche quality filter^43^ was applied. Reads from all samples were co-assembled using megahit ^44^. We then used the anvi’o v7.1^45^ metagenomic workflow to compute coverage for each gene across metagenomes, and to refine metagenome-assembled-genomes (MAGs). Briefly, the workflow uses (1) Prodigal v2.6.3^46^ to identify open-reading frames (ORFs), (2) *’anvi-run-hmm*’ to identify single copy core genes from bacteria (n=71) and ribosomal RNAs (n=12) using HMMER v3.3^47^, (3) ’*anvi-run-pfams*’, ’*anvi-run-kegg-kofams*’, and ’*anvi-run-cazymes’* to annotate ORFs with EBI’s PFAM database^48^, the KOfam HMM database of KEGG orthologs (KOs)^49^, and the dbCAN CAZyme HMM database^50^, respectively. We used Bowtie2 v2.3.5.1^51^ to recruit metagenomic short-reads to the contigs, and samtools v1.11^52^ to convert SAM files to BAM files. We profiled the resulting BAM files with ’*anvi-profile’* and used the program ’*anvi-merge’* to combine all single profiles into a merged profile for downstream visualization. We used *’anvi-export-gene-coverage-and-detection*’ to generate coverage tables for downstream analysis in R. ’*deseq2*’ was used on the exported count data to identify differentially abundant KOs across time points and treatment groups. To identify CAZyme substrate utilization functions, EC numbers from KEGG annotations were used to map to the dbCAN-sub database^50^. All plots were generated with the R package ’*tidyverse*’^53^ or GraphPad Prism (GraphPad Software).

### Metabolite Extraction from Fecal/Cecal Material

Metabolites were extracted with the addition of extraction solvent (80% methanol spiked with internal standards and stored at -80°C, Table S5) to pre-weighed fecal/cecal samples at a ratio of 100 mg of material per mL of extraction solvent in beadruptor tubes (Fisherbrand; 15-340-154). Samples were homogenized at 4°C on a Bead Mill 24 Homogenizer (Fisher; 15-340-163), set at 1.6 m/s with 6 thirty-second cycles, 5 seconds off per cycle. Samples were then centrifuged at - 10°C, 20,000 x g for 15 min and the supernatant was used for subsequent metabolomic analysis.

Cecal samples were used in lieu of fecal samples for SCFA analysis as the cecum is the primary site of SCFA production via fermentation in the gut^54^.

### Metabolite Analysis using GC-EI-MS and Methoxyamine and TMS Derivatization

Metabolites were analyzed using gas chromatography mass spectrometry (GCMS) with electron impact ionization. To a mass spectrometry autosampler vial (Microliter; 09-1200), 100 µL of metabolite extract was added and dried down completely under a nitrogen stream at 30 L/min (top) and 1 L/min (bottom) at 30°C (Biotage SPE Dry 96 Dual; 3579M). To dried samples, 50 µL of freshly prepared 20 mg/mL methoxyamine (Sigma; 226904) in pyridine (Sigma; 270970) was added and incubated in a thermomixer C (Eppendorf) for 90 min at 30°C and 1400 rpm.

After samples were cooled to room temperature, 80 µL of derivatizing reagent (BSTFA + 1% TMCS; Sigma; B-023) and 70 µL of ethyl acetate (Sigma; 439169) were added and samples were incubated in a thermomixer at 70°C for 1 hour and 1400 rpm. Samples were cooled to RT and 400 µL of Ethyl Acetate was added to dilute samples. Turbid samples were transferred to microcentrifuge tubes and centrifuged at 4°C, 20,000 x g for 15 min. Supernatants were then added to mass spec vials for GCMS analysis. Samples were analyzed using a GC-MS (Agilent 7890A GC system, Agilent 5975C MS detector) operating in electron impact ionization mode, using a HP-5MSUI column (30 m x 0.25 mm, 0.25 µm; Agilent Technologies 19091S-433UI) and 1 µL injection. Oven ramp parameters: 1 min hold at 60°C, 16°C per min up to 300°C with a 7 min hold at 300°C. Inlet temperature was 280°C and transfer line was 300°C. Data analysis was performed using MassHunter Quantitative Analysis software (version B.10, Agilent Technologies) and confirmed by comparison to authentic standards. Normalized peak areas were calculated by dividing raw peak areas of targeted analytes by averaged raw peak areas of internal standards. Bile acid assays for cholic acid also included allocholic acid; assays for lithocholic acid also included allolithocholic acid and isolithocholic acid.

### Histopathology

Cecal tissue cross-sections were fixed with 4% formalin for 24 hours and were stored in 70% ethanol until paraffin embedding and tissue sectioning. Embedding, sectioning, and H&E staining were performed by the University of Chicago Human Tissue Resource Center. Each tissue section was scored for pathology in a blinded fashion by a pathologist and a trained researcher according to the system outlined by Barthel, *et al*.^16^ Briefly, two independent scores for submucosal edema, PMN infiltration, goblet cells, and epithelial integrity were averaged for each tissue sample. The combined histopathological score for each sample was determined as the sum of these averaged scores. It ranges between 0 and 12 arbitrary units and covers the following levels of inflammation: 0 = no signs of inflammation; 1-2 = minimal signs of inflammation; 3-4 = slight inflammation; 5-8 = moderate inflammation; 9-13 = profound inflammation.

### RT-qPCR

Total messenger RNA isolated from colonic mucosal scrapings was used with Transcriptor First Strand cDNA Synthesis Kit (Roche Diagnostics Corporation) to obtain cDNA. Real-time qPCR was performed using iTaq Universal SYBR Green Supermix with CFX384 Real-Time System (Bio-Rad). Primers and cycling conditions were derived from Devlin et al., 2022^55^ (Table S10). Expression was calculated via ΔΔCt relative to the housekeeping gene *actb* and the control group RC-PBS-PBS.

### Construction of Probabilistic Annotation from ASV Sequences

We mapped as many of the ASV sequences as possible from our amplicon sequencing and analysis pipeline to 16S sequences of full reference genomes that are a part of the AGORA2 set of common gut microorganisms^56^. We searched for identical 16S sequences and iteratively reduced the identity threshold by 1% to 90% until a match was acquired, or accepted that the ASV does not have a matching AGORA2 genome if no match was acquired at the 90% threshold. We mapped 3035 distinct AGORA2 reference genomes to 1654 of the experimentally detected ASVs, and then created 267 ASVsets by grouping ASVs whose reference genomes were all more than 50% identical with each other. The ASVsets were named for the most common genus of each set and appended with an iterative suffix if the genus had already been identified by a previous ASVset (e.g. Lactoccocus.1, then Lactoccous.2, *etc*), which indicates that the ASVsets embody sub-genus phylogenetic resolution. All mapped reference genomes were loaded into KBase^57^, annotated with RAST^58^ (https://narrative.kbase.us/narrative/178418), and then the reference genomes were merged into a single probabilistic annotation for each ASVset that contains a pseudo function-based gene representing each distinct RAST-assigned function annotated across all reference genomes mapped to the ASVset. A probability *p*_*func*_ for each pseudogene in the probabilistic annotation was created from the count of the associated function across the reference genomes of the ASVset divided by the number of reference genomes in the ASVset. These probabilistic annotations were saved in KBase as genome objects, with a name matching the name of the ASVset from which they were derived (see https://narrative.kbase.us/narrative/181152). Each pseudogene was annotated with the corresponding RAST function and the associated probability saved in the evidence score. In the alias list for each pseudogene, we included the full list of gene IDs annotated with the associated RAST function across all the reference genomes comprising the probabilistic annotation.

### Reconstruction of ASVset prGEMs Based on Probabilistic Annotations

We applied the MS2 - Build Prokaryotic Metabolic Models with OMEGGA app in KBase^1^ to reconstruct a draft metabolic model from the probabilistic annotation for each ASVset. Reactions are mapped through this reconstruction process to the pseudo function-based genes comprising each probabilistic annotation. The draft models were gapfilled in glucose minimal media, to ensure that every model includes all the reactions needed to permit growth without any essential auxotrophy, and were further gapfilled on all of the 63 metabolites, to ensure that the models were capable of consmption, production, and growth in metabolic environment determined from our metabolomics data. The gapfilling ensures that all of the pathways needed to interact with the measured metabolites were available and ensures that models can function in diverse contexts that the microbiome environment presents; however, we are not asserting that all ASVs have all of these gapfilled capabilities. All reactions in these ASVset probabilistic genome-scale models (prGEM) are assigned a probability *p*_*rxn*_ based on their pseudogene associations. Reactions with pseudogenes are assigned the highest *p*_*func*_ associated with any of the pseudogenes to which the reaction was matched (see the previous section for how pseudogene probabilities are computed), while gapfilled reactions are assigned a probability of 0. The prGEMs were saved in KBase using the same name as their associated ASVset: https://narrative.kbase.us/narrative/181152.

### Generation of Strain-Metabolite Interaction Probability Profiles (SMIPPs)

To gain insights into the metabolic potential of our ASVsets based on their probabilistic annotations, we applied flux balance analysis (FBA) to compute the probability that each ASVset will interact with each of the 63 metabolites measured in our experimental samples. We simulated three possible interactions for each metabolite: (1) consumption and transformation by forcing a negative exchange flux; (2) production, after conversion from a different input nutrient, by forcing a positive exchange flux; and (3) growth by forcing biomass production in a minimal media with the metabolite as the sole carbon source. The forced conditions described above were achieved through tailored FBA constraints and an FBA objective that minimized the product of each forward or reverse reaction flux and the probability that the reaction should not be part of the model: 1 − *p*_*rxn,i*_ for reaction *i*. This objective function determines the most likely reactions and pathway for each phenotype, which enables computing an overall probability for each phenotype by averaging the *p*_*rxn,i*_ values associated with all the reactions involved in the pathway. Based on this analysis, we created three species-metabolite interaction probability profiles (SMIPPs) across our ASVsets: consumption, production, and growth. These SMIPPs take the form of matrices – *P*_*SMIPP*,*up*_, *P*_*SMIPP*,*ex*_, and *P*_*SMIPP*,*gr*_ for production, consumption, and growth, respectively – where the rows correspond to ASVsets, the columns correspond to metabolites, and the data elements contain the average probability of the reactions used in the associated ASVset prGEM to implement a given phenotype (consumption, production, or growth) with the associated metabolite. We used cluster heatmaps to render the *P*_*SMIPP*,*up*_, *P*_*SMIPP*,*ex*_, and *P*_*SMIPP*,*gr*_ matrices, grouping metabolites and ASVsets that had similar profiles in Figure S5.

### Generation of Microbiome-Metabolite Interaction Probability Profiles (MMIPPs)

Combining SMIPPs with microbiome composition data allowed us to translate strain interactions of each SMIPP matrix into sample-wide aggregate microbiome interactions with the same metabolites. The sample ASV abundance matrix, *A*_*ASV*_, was first translated into an ASVset abundance matrix, *A*_*set*_ , to align with the SMIPPs that were only available for ASVsets. *A*_*ASV*_ has dimensions of *a*_*strain*_ (number of ASVs) by *s* (number of samples), with each element being the ASVset abundance in given a sample. Each ASVset abundance of *A*_*set*_ is computed by summing the rows of *A*_*ASV*_for all members of the ASVset. The dot product of *A*_*set*_ and each PSMIPP matrix (*P*_*SMIPP*,*up*_, *P*_*SMIPP*,*ex*_, and *P*_*SMIPP*,*gr*_ ) were computed to produce new corresponding *P*_*SMIPP*,*up*_, *P*_*SMIPP*,*ex*_, and *P*_*SMIPP*,*gr*_ matrices, which are *s* (# samples) x *m* (# metabolomics metabolites) matrices with the probabilities that the microbiome in each sample has the metabolic potential to consume, produce, or grow on each measured metabolite.

### Correlating Metabolite Interaction Potential with Metabolite Abundance

The *P*_*SMIPP*_ matrices importantly represent the metabolic capacity for an interaction, and not the probability of that interaction. Correlating *P*_*SMIPP*_ with the metabolite abundance matrix *M*, an *s* x *m* matrix with elements of abundance of each metabolite in each sample, therefore identifies which metabolites might be most impacted by microbial metabolism. Metabolite correlations in Supplementary Table S8 were determined by correlating the probabilities of *P*_*SMIPP*,*up*_, *P*_*SMIPP*,*ex*_, and *P*_*SMIPP*,*gr*_ with the abundances in *M.* Highly positively or negatively correlated metabolites signify those that are most likely affected by microbial metabolism.

### Construction of Community Metabolic Models for Sample Intervals

We created a community prGEM of each interval between experimental samples for which metabolomics data was collected using the compartmentalized community model formalism described in Henry *et al.,* 2016^59^. The *A*_*int*,*set*_ matrix of ASVset abundances per sample interval is derived from the average of *A*_*set*_ between adjacent columns (i.e. consecutive timepoints) that defined the interval bounds. Interval community models were built for each column in *A*_*int*,*set*_ by merging the ASVset prGEMs with relative abundances > 1% into a single probabilistic community model (prcGEM). Each ASVset prGEM compartment was assigned an index, which was appended to the IDs of the transport reactions and the intracellular metabolites and reaction IDs of the contained prGEM, while exchange fluxes and extracellular metabolite IDs were shared by all prGEM and thus were not indexed. Finally, the objective function for each prcGEM was defined as one gram of community biomass being the dot product of the biomass flux and ASVset abundance from the *A*_*int*,*set*_ among all merged prGEMs in the interval. The prcGEM therefore contains the union of ASVsets between the samples that define the interval, which allows the model to capture potential ASVsets interactions between the first and second samples of the interval. All of these interval prcGEMs were saved in KBase: https://narrative.kbase.us/narrative/181152.

### Simulation of Interval Community Models to Predict Maximum Likelihood Interactions Between ASVsets and Metabolites

FBA simulations of each interval prcGEM predicted the maximum-likelihood interactions between the ASVsets and the measured metabolites in each sample interval. Constraints were adjusted to simulate rich media that includes all compounds that can be utilized by any ASVset in the prcGEM, which reflects both uncertainty in and the complexity of the nutritional environment of the gut microbiome. An upper limit of 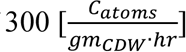 was placed on the total carbon uptake of the prcGEM to reflect nutrient limitations, thereby forcing resource competition and incentivizing the consumption of the most nutritious resources. All dipeptide exchanges were forced to zero because they add little value for understanding microbiome behavior yet combinatorially increase complexity compared to simple amino acid exchanges. A flux capacity constraint was added to that limited net flux through each ASVset compartment to less than 750 times the growth rate of the compartment, which permits non-auxotrophic optimal growth on glucose in E. coli but prevents an ASV from carrying far more flux than is justified by its level of abundance within the microbiome. Oxygen uptake was set to 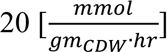, which means that the microbiome is operating at one-sixth of the oxygen required for fully aerobic growth combined with the specified carbon uptake limit of 300. The community biomass reaction, with all of the aforementioned constraints, was then maximized and constrained to be at least 50% of this optimal growth to force effective community growth while permitting other specifications of the probabilistic optimization to be met. Constraints on the exchange fluxes for metabolites observed in our metabolomics data were then added, such that the exchange flux of metabolites with increasing or decreasing concentration over an interval were forced to be positive or negative, respectively, thereby simulating production or consumption of the metabolite by the microbiome. The fluxes of these concentration-based constraints were increased until just before the optimization became infeasible to optimally replicate the observed metabolite trajectories in the community simulations. Finally, the model objective was defined to minimize the dot product of each reaction flux and the inverse probability (1 − *P*_*rxn*_) of the associated prGEM for all ASVset prGEMs to acquire the maximum-likelihood fluxes considering the metabolic capacity and abundance of each individual ASVset for community growth and to reproduce the observed metabolite trajectories.

The resultant community fluxes from simulating these communities were visualized in the Escher Map in Figure 3 that depicts metabolic interactions among the ASVsets through the studied ecological successions after antibiotics treatment.

### Code availability

The Jupyter Notebooks in which the modeling data was processed and the figures were developed is accessible at https://github.com/HenryLabResearch/ABX_mouse_gut

### Data availability

The data, including all DNA sequencing datasets, that support the findings of this study are available in this article, the Supplemental Information, and BioProject accession PRJNA992061.

## ACKNOWLEDGMENTS

We thank the Chang lab members for scientific support received. This work was performed with support from NIH T32DK007074 (M.S.K.), NIH RC2DK122394 (E.B.C.), NIH T32GM007281 (M.S.K.), InnoHK via the Hong Kong Innovation and Technology Commission, the Host-Microbe and Tissue and Cell Engineering cores of the UChicago DDRCC, Center for Interdisciplinary Study of Inflammatory Intestinal Disorders (C-IID) - (NIDDK P30 DK042086), the Gastrointestinal Research Foundation of Chicago, and The Simons Foundation (J.B.). C.S.H., A.F., and K.B. were supported by the KBase project of the U.S. Department of Energy, Office of Science, Office of Biological and Environmental Research (DE-AC02-06CH11357).

## AUTHOR CONTRIBUTIONS

Conceptualization, M.S.K., E.B.C., J.B.; Methodology, M.S.K., E.B.C., J.B., A.F., K.B., C.S.H.; Formal Analysis, M.S.K., A.F., K.B., C.S.H., A.G.; Investigation, M.S.K., M.C., M.L.S., M.K., C.C.; Data Curation, M.S.K., K.B., A.F.; Writing – Original Draft, M.S.K; Writing – Review and Editing, M.S.K., A.F., A.G., M.L.S., C.S.H., E.B.C., S.C.N., F.C., D.R., J.B.; Visualization, M.S.K., A.F.; Funding Acquisition, C.S.H., J.B., S.C.N., F.C., E.B.C.

## ADDITIONAL INFORMATION

Supplementary Information is available for this paper.

Correspondence and requests for materials should be addressed to E. Chang.

## EXTENDED DATA FIGURES

**Figure S1:**
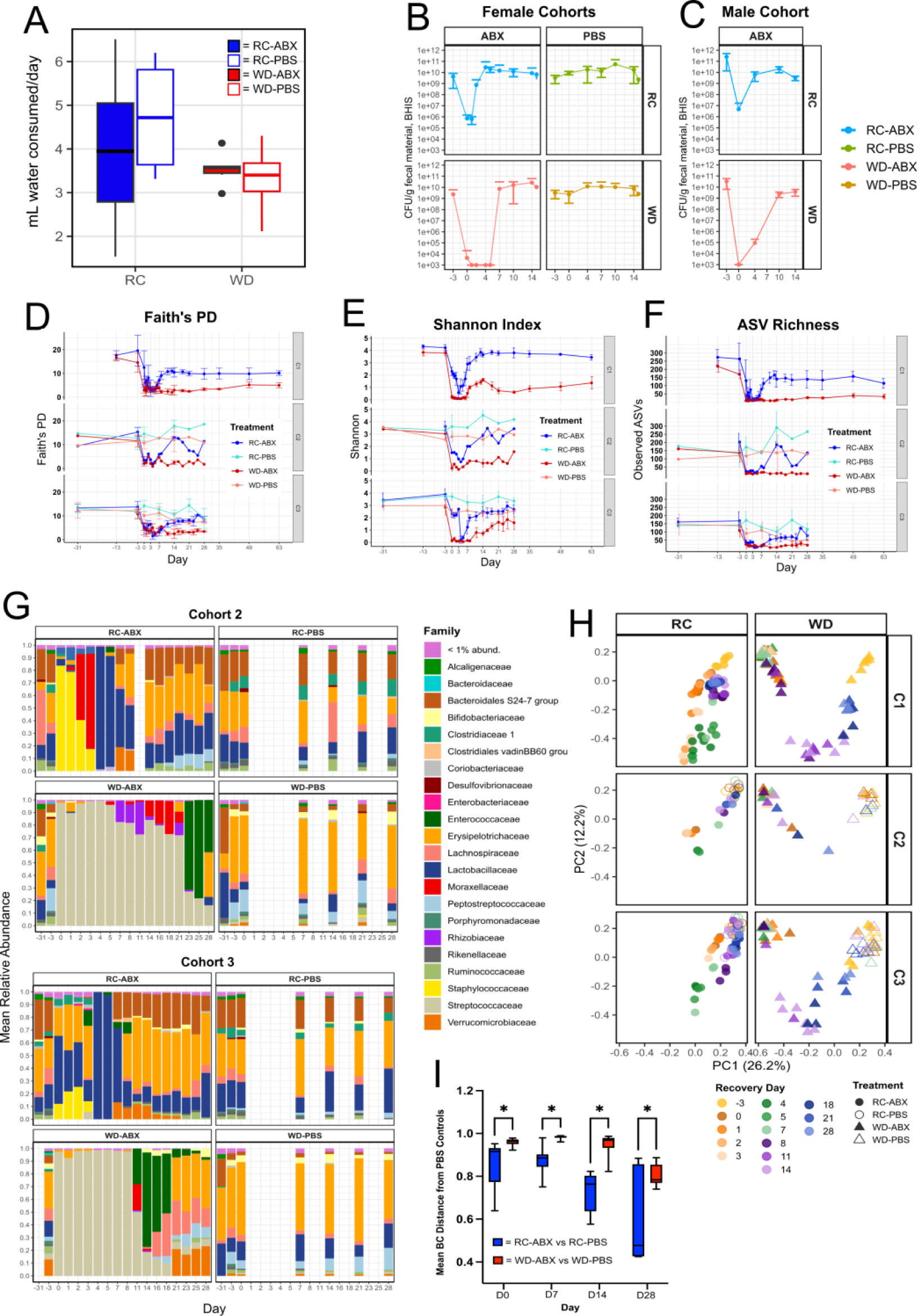
Western diet impairs microbiome taxonomic and biomass recovery from antibiotics. (A) Consumption of ABX- or PBS-spiked water per mouse per day did not differ significantly across any treatment groups (n=4-6/group, one-way ANOVA). (B-C) Microbial CFUs plated on anaerobic BHIS media from all (B) female (n=10-24/group) and (C) male cohorts (n=6/group) through Day 14 of recovery post-ABX. Three of six female cohorts and the male cohort did not undergo 16S analysis as in the rest of Figure 1; these data are therefore excluded from Figure 1A and Table S1A, but are analyzed separately in Table S1B. (D-F) Comparison of alpha diversity metrics across cohorts over time. (D) Faith’s phylogenetic diversity; (E) Shannon index; (F) ASV richness. Statistics are presented in Table S2. (G) Mean relative abundances of different microbial families for Cohorts 2 and 3. (H) PCoA of 16S-based microbiome taxonomic composition at the genus level using Bray-Curtis dissimilarity for samples from all treatment groups and cohorts through Day 28 of recovery. (I) Mean Bray-Curtis dissimilarity of antibiotic-treated groups from their respective PBS control groups at each timepoint (Table S2).

**Figure S2:**
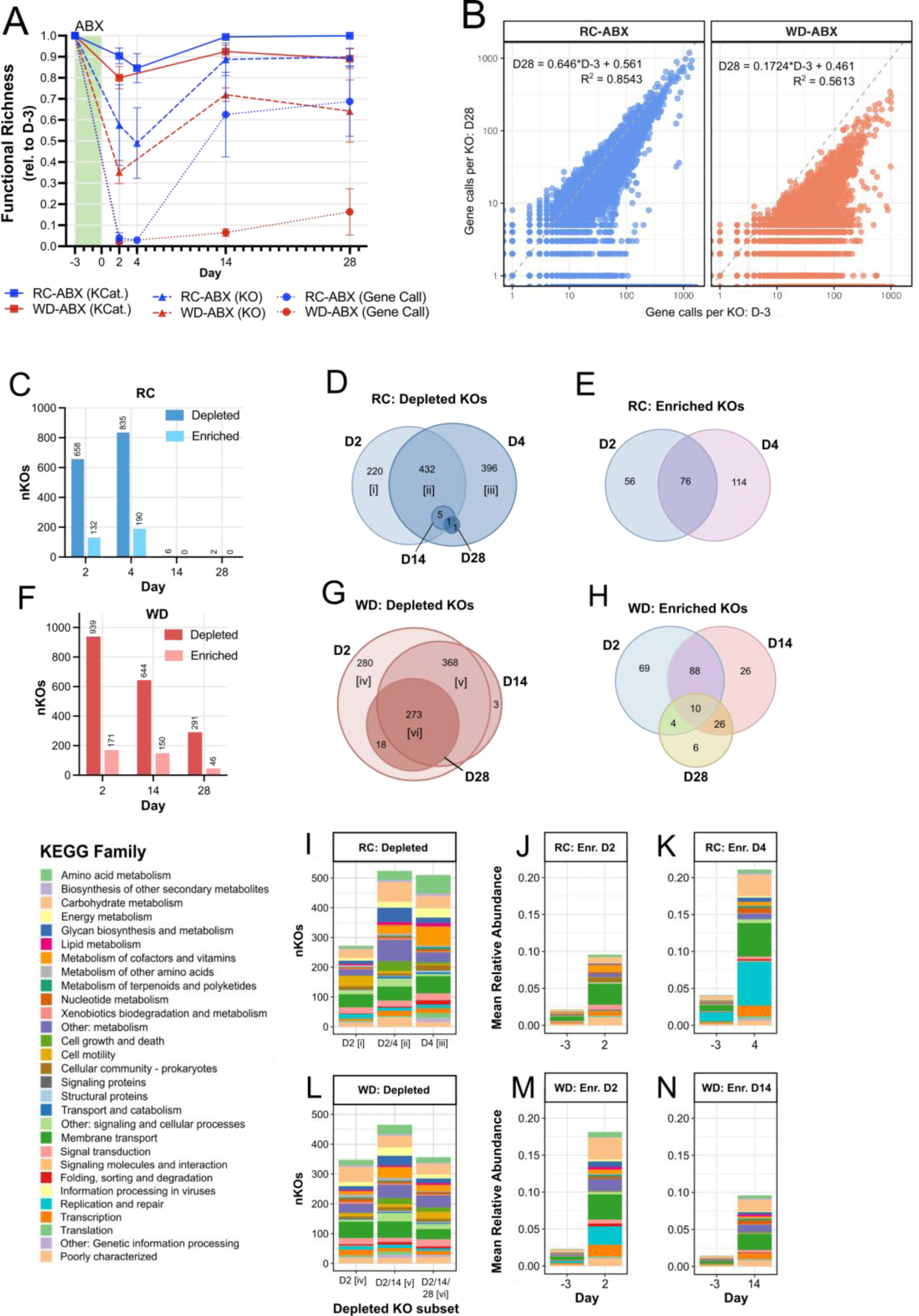
Microbiome metagenomic recovery dynamics differ across dietary treatments. (A) Metagenomic functional richness in fecal samples from mice on RC-ABX (blue) and WD-ABX (red) at the KEGG Category (KCat), KEGG Ortholog (KO), and gene call level as a percentage of functional richness at Day -3 (pre-ABX) (n=3/group, Table S3). (B) Initial (Day - 3) versus final (Day 28) functional redundancy (genes calls per KO) for mice on RC-ABX (blue) and WD-ABX (red) (Table S3). For mice on RC-ABX (C-E) or WD-ABX (F-H), counts of significantly differentially abundant KOs (C, F), and Venn diagrams of depleted (D, G) or enriched (E, H) KOs across timepoints. KEGG Family mapping of significantly depleted KOs in mice on (I) RC-ABX or (L) WD-ABX. Roman numerals indicate the subset of KOs depicted in panels (D) and (G). Relative abundances of significantly enriched KOs in mice on RC-ABX at (J) Day 2 and (K) Day 4 relative to Day -3. Relative abundances of significantly enriched KOs in mice on WD-ABX at (M) Day 2 and (N) Day 14 relative to Day -3. See Table S4 for statistics.

**Figure S3:**
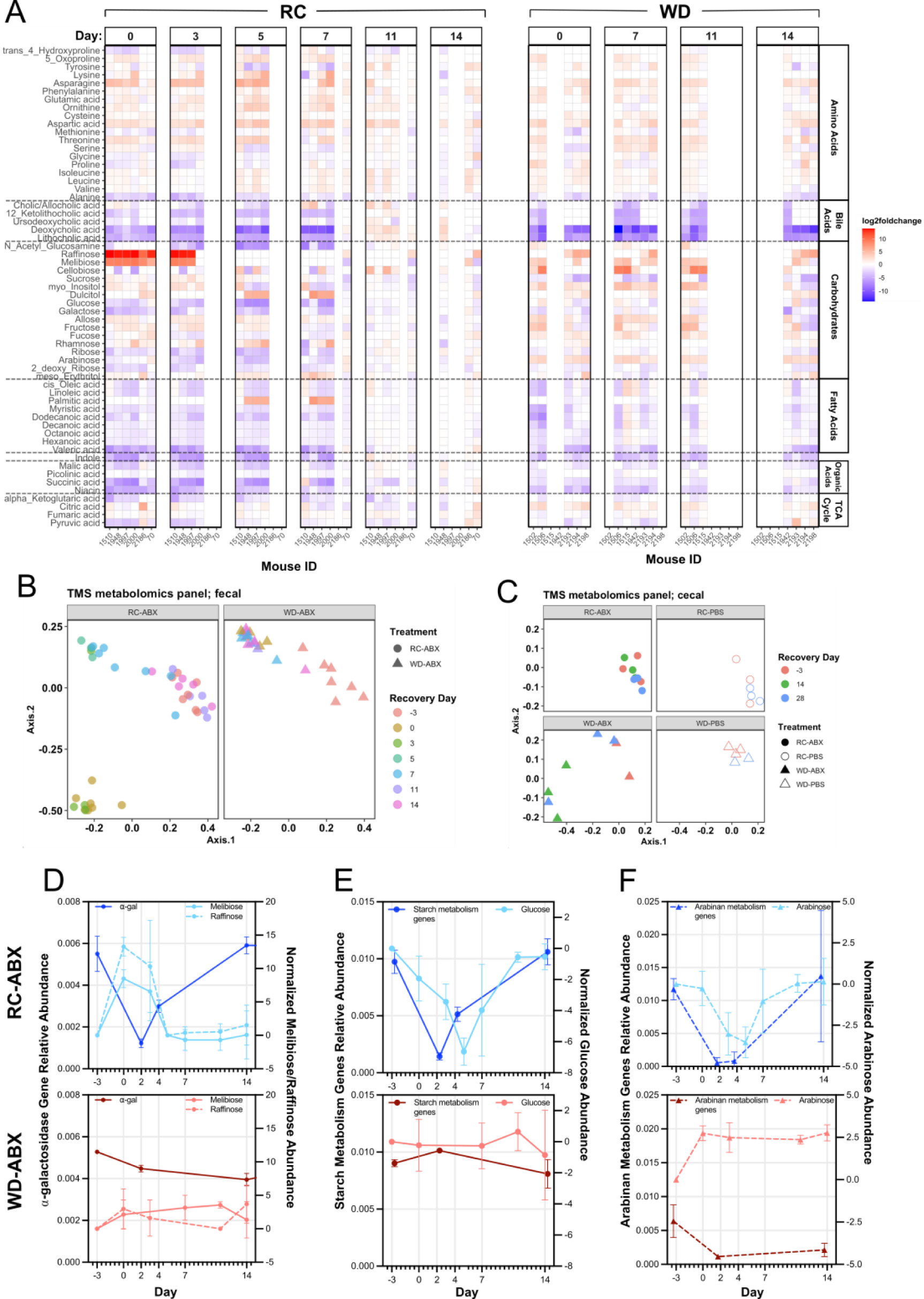
Metabolomic evaluations show distinct recovery dynamics across diets. (A) Normalized metabolite abundances for mice on RC-ABX at different timepoints are consistent across mice. Each vertical block represents a different day of recovery. Each column within a block represents samples from a different mouse. Abundances are normalized to Day -3 (pre-ABX) for each mouse. (B) PCoA of fecal metabolomics TMS panel data using Bray-Curtis dissimilarity for samples from RC-ABX and WD-ABX through Day 14 of recovery. (C) PCoA of cecal metabolomics TMS panel data using Bray-Curtis dissimilarity for samples from RC- ABX and WD-ABX through Day 28 of recovery. Cecal samples were used due to availability of material through Day 28. (D – F) Metagenomic gene abundances (left axis, Materials and Methods) and normalized metabolite abundances (right axis) over time for mice on RC (top, blue) and WD (bottom, red). (E) ⍺-galactosidase genes, melibiose and raffinose abundance. (F) Starch metabolism genes, glucose abundance. (G) Arabinan metabolism genes, arabinose abundance. See Table S6 for statistics.

**Figure S4:**
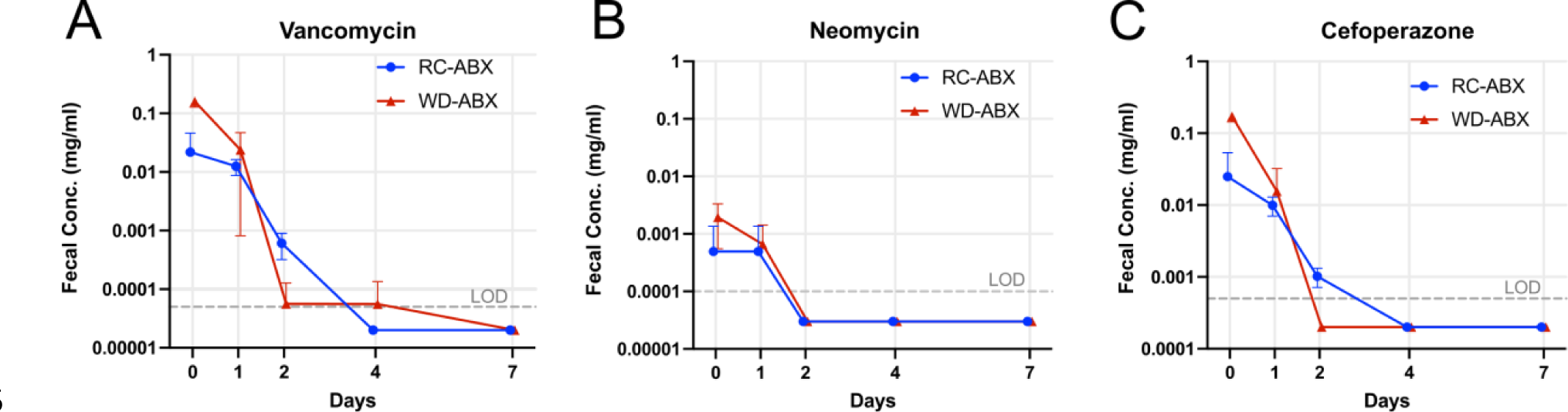
Residual antibiotic concentrations were not significantly different across RC- ABX and WD-ABX groups. Absolute quantification of fecal (A) vancomycin, (B) neomycin, and (C) cefoperazone from immediately after cessation fo antibiotic treatment through Day 7 of recovery. See Table S7 for statistics.

**Figure S5:**
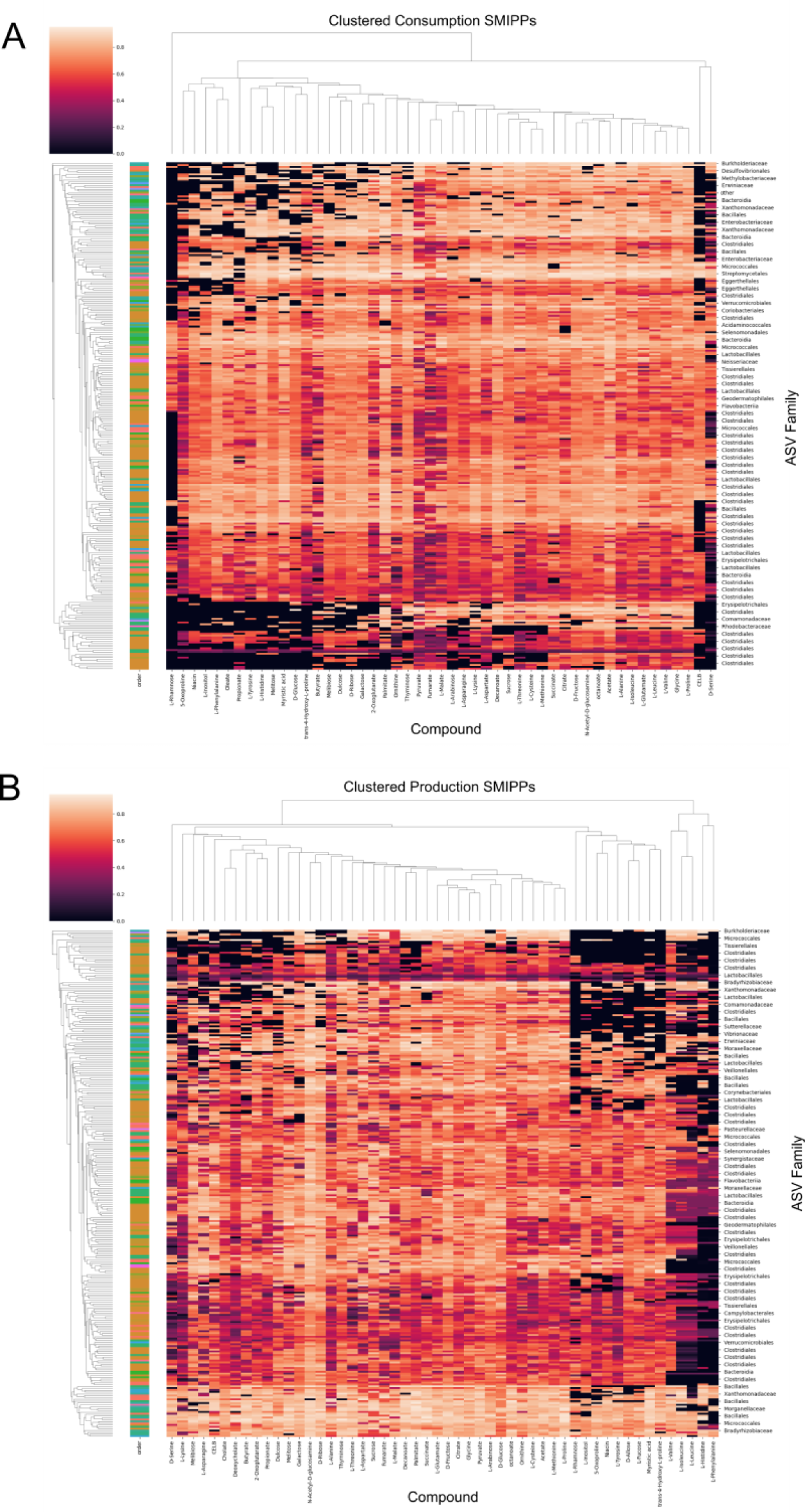
Strain-metabolite Interaction Probability Profiles (SMIPPs) reveal metabolic specialization. Heatmaps indicating the probability that a given ASV prGEM (rows) has the capacity to (A) consume or (B) produce the indicated compounds (columns).

**Figure S6:**
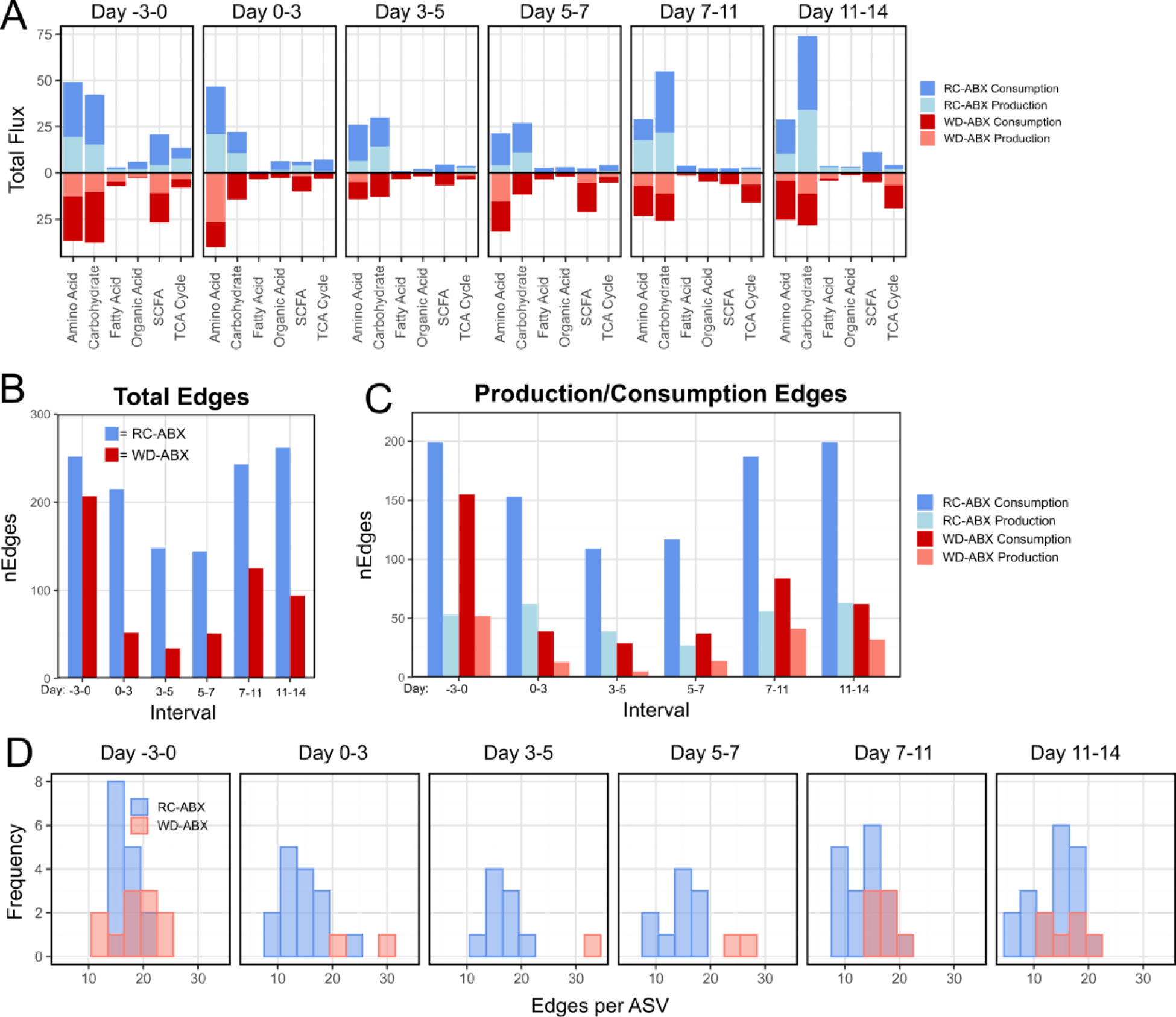
Network metrics of community flux simulations vary across dietary treatment groups. (A) Total predicted consumption or production flux through each metabolite category in mice on RC (blue, top) or WD (red, bottom) over the indicated recovery interval. As recovery proceeds, mice on RC push more flux through carbohydrate metabolism than mice on WD. (B) Total edges (i.e. metabolic interactions) in the community flux-balance analysis simulation networks across dietary groups at each time interval, broken down by (C) production or consumption edges. The microbiome of mice on RC has more edges at all timepoints, indicating that they have more/broader metabolite interaction (primarily consumption interactions) than in mice on WD. (D) Histograms depicting the distribution of edges per ASV across diet groups at each time interval. Mice on WD have few taxa that interact with a large number of metabolites, whereas in mice on RC, a broader array of taxa interact with an intermediate number of metabolites.

**Figure S7:**
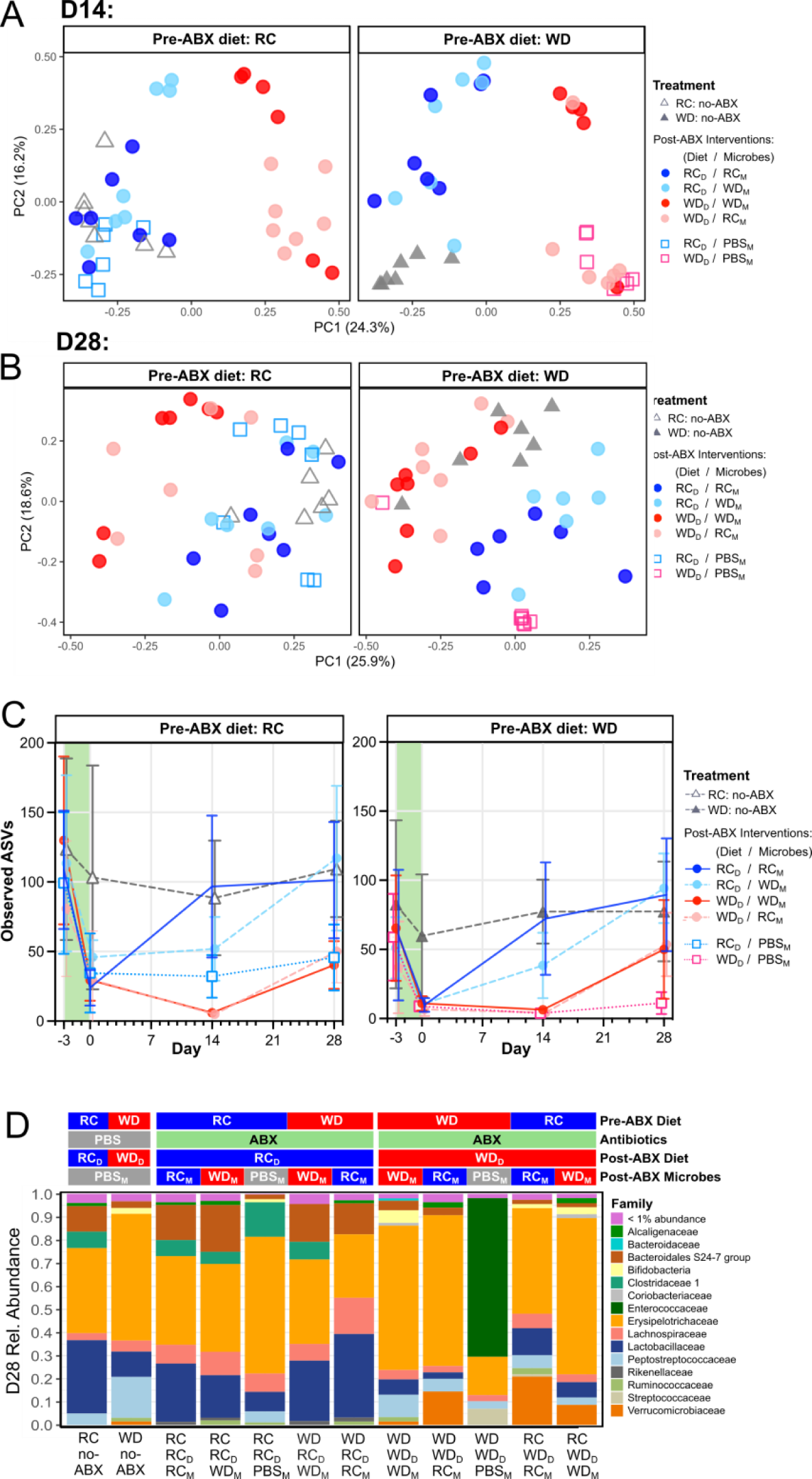
Dietary intervention and microbial transplant effects through Day 28 of recovery. PCoA plot of 16S-based taxonomic data for mice on all treatment groups at D14 (A) and D28 (B) of recovery. Data is paneled according to pre-ABX diet. (C) ASV richness of all treatment groups through Day 28 of recovery. Data is paneled according to pre-ABX diet. (D) Mean relative abundances of microbial families at Day 28 across treatment groups.

**Figure S8:**
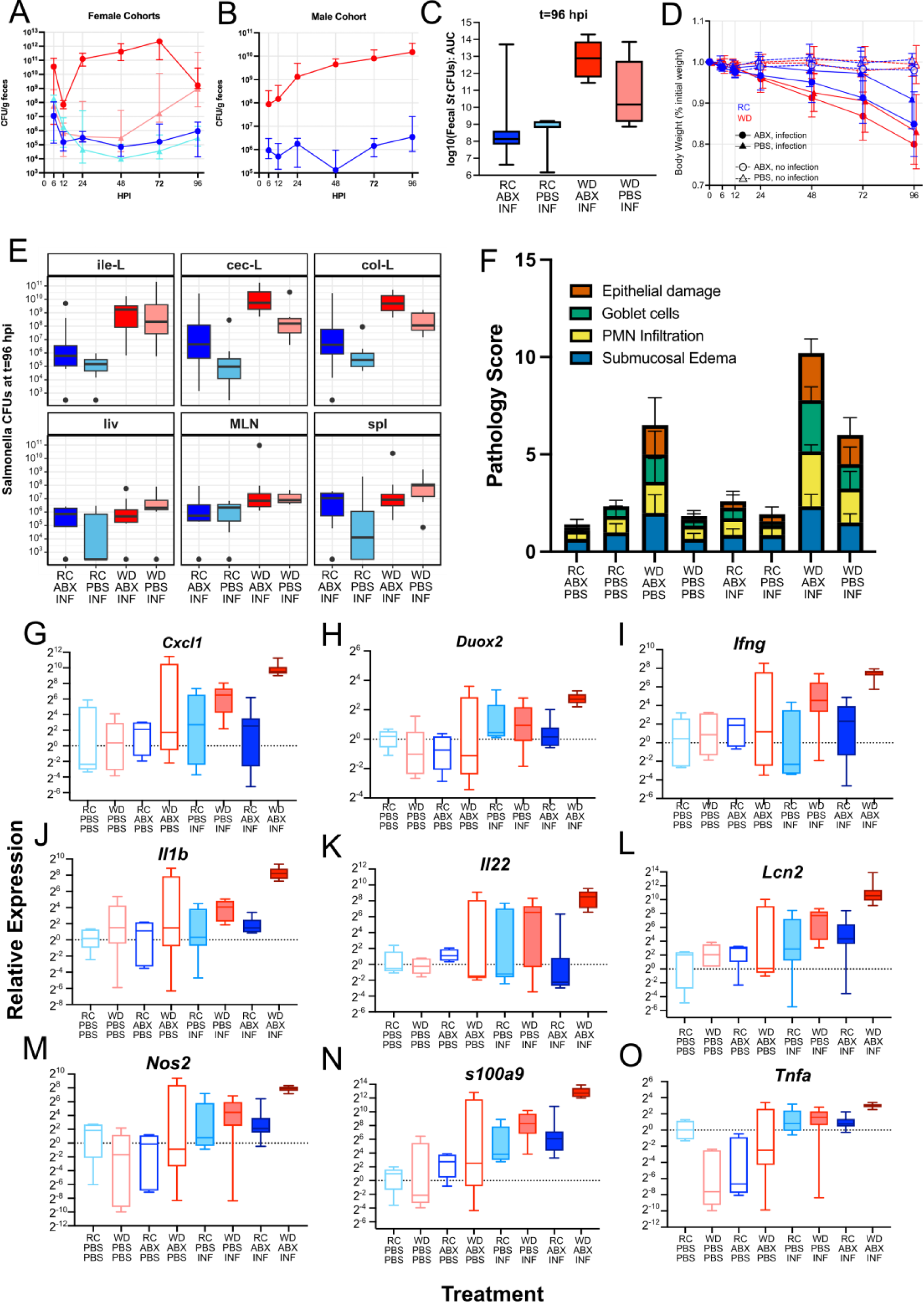
Supplemental information regarding colonization resistance experiments. ***St*** CFU counts from female (A) and male (B) cohorts through t=96 hpi. (C) Log10 transformed Infection AUC for all infected treatment groups. (D) Body weight after infection as a percentage of pre-infection body weight for all treatment groups. (E) *St* CFU counts across body tissue sites for all infected treatment groups at t=96 hpi. (F) Cecal and (G) colonic histopathology scoring of all treatment groups at t=96 hpi broken down by subscore. (H-P) mRNA expression of immune genes in cecal mucosal scrapings at t=96 hpi based on RT-qPCR. Expression is normalized to the housekeeping gene Actb and the RC-PBS-PBS treatment group. See Table S10 for statistics and additional information.

## REFERENCES

1. Bisanz, J. E., Upadhyay, V., Turnbaugh, J. A., Ly, K. & Turnbaugh, P. J. Meta-Analysis Reveals Reproducible Gut Microbiome Alterations in Response to a High-Fat Diet. Cell Host & Microbe 26, 265–272.e4 (2019).

2. Sonnenburg, E. D. & Sonnenburg, J. L. Starving our Microbial Self: The Deleterious Consequences of a Diet Deficient in Microbiota-Accessible Carbohydrates. Cell Metabolism 20, 779–786 (2014).

3. Nieuwdorp, M., Gilijamse, P. W., Pai, N. & Kaplan, L. M. Role of the Microbiome in Energy Regulation and Metabolism. Gastroenterology 146, 1525–1533 (2014).

4. Asnicar, F. et al. Microbiome connections with host metabolism and habitual diet from 1,098 deeply phenotyped individuals. Nat Med 27, 321–332 (2021).

5. Turnbaugh, P. J. et al. An obesity-associated gut microbiome with increased capacity for energy harvest. Nature 444, 1027–131 (2006).

6. Levy, M., Kolodziejczyk, A. A., Thaiss, C. A. & Elinav, E. Dysbiosis and the immune system. Nature Reviews Immunology 2017g 17:4 17, 219–232 (2017).

7. Ng, K. M. et al. Recovery of the Gut Microbiota after Antibiotics Depends on Host Diet, Community Context, and Environmental Reservoirs. Cell Host and Microbe 26, 650–665.e4 (2019).

8. Tanes, C. et al. Role of dietary fiber in the recovery of the human gut microbiome and its metabolome. Cell Host & Microbe 29, 394–407.e5 (2021).

9. Taur, Y. et al. Reconstitution of the gut microbiota of antibiotic-treated patients by autologous fecal microbiota transplant. Sci Transl Med 10, eaap9489 (2018).

10. Tropini, C. et al. Transient Osmotic Perturbation Causes Long-Term Alteration to the Gut Microbiota. Cell 173, 1742–1754.e17 (2018).

11. Shade, A. et al. Fundamentals of Microbial Community Resistance and Resilience. Frontiers in Microbiology 3, (2012).

12. May, R. M. Will a Large Complex System be Stable? Nature 238, 413–414 (1972).

13. Miyoshi, J. et al. Minimizing confounders and increasing data quality in murine models for studies of the gut microbiome. PeerJ 6, e5166 (2018).

14. Kennedy, M. S. et al. Dynamic genetic adaptation of Bacteroides thetaiotaomicron during murine gut colonization. Cell Rep 42, 113009 (2023).

15. Buffie, C. G. & Pamer, E. G. Microbiota-mediated colonization resistance against intestinal pathogens. Nature Reviews Immunology 13, 790–801 (2013).

16. Barthel, M. et al. Pretreatment of Mice with Streptomycin Provides a Salmonella enterica Serovar Typhimurium Colitis Model That Allows Analysis of Both Pathogen and Host. Infection and Immunity 71, 2839 (2003).

17. Wotzka, S. Y. et al. Escherichia coli limits Salmonella Typhimurium infections after diet shifts and fat-mediated microbiota perturbation in mice. Nat Microbiol 4, 2164–2174 (2019).

18. Santos, R. L. et al. Animal models of Salmonella infections: enteritis versus typhoid fever. Microbes and Infection 3, 1335–1344 (2001).

19. NHANES. NHANES - What We Eat in America: Nutrient Intakes from Food by Gender and Age. vol. National Health and Nutrition Examination Survey Data. (U.S. Department of Health and Human Services, Centers for Disease Control and Prevention, Hyattsville, MD, 2009).

20. Wang, J., Wang, P., Wang, X., Zheng, Y. & Xiao, Y. Use and Prescription of Antibiotics in Primary Health Care Settings in China. JAMA Internal Medicine 174, 1914–1920 (2014).

21. Bell, M. Antibiotic Misuse: A Global Crisis. JAMA Internal Medicine 174, 1920–1921 (2014).

22. Dethlefsen, L. & Relman, D. A. Incomplete recovery and individualized responses of the human distal gut microbiota to repeated antibiotic perturbation. Proceedings of the National Academy of Sciences of the United States of America 108, 4554–4561 (2011).

23. Shaw, L. P. et al. Modelling microbiome recovery after antibiotics using a stability landscape framework. The ISME Journal 13, 1845–1856 (2019).

24. Zaura, E. et al. Same Exposure but two radically different responses to antibiotics: Resilience of the salivary microbiome versus long-term microbial shifts in feces. mBio 6, (2015).

25. Palleja, A. et al. Recovery of gut microbiota of healthy adults following antibiotic exposure. Nature Microbiology 3, 1255–1265 (2018).

26. Chng, K. R. et al. Metagenome-wide association analysis identifies microbial determinants of post- antibiotic ecological recovery in the gut. Nature Ecology and Evolution 4, 1256–1267 (2020).

27. Ledder, O. & Turner, D. Antibiotics in IBD: Still a Role in the Biological Era? Inflammatory Bowel Diseases 24, 1676–1688 (2018).

28. Jowett, S. L. et al. Influence of dietary factors on the clinical course of ulcerative colitis: a prospective cohort study. Gut 53, 1479–1484 (2004).

29. Cohen, N. A. & Maharshak, N. Novel Indications for Fecal Microbial Transplantation: Update and Review of the Literature. Dig Dis Sci 62, 1131–1145 (2017).

30. Ritchie, M. L. & Romanuk, T. N. A Meta-Analysis of Probiotic Efficacy for Gastrointestinal Diseases. PLoS ONE 7, e34938 (2012).

31. O’Toole, P. W., Marchesi, J. R. & Hill, C. Next-generation probiotics: the spectrum from probiotics to live biotherapeutics. Nat Microbiol 2, 1–6 (2017).

32. Gilbert, J. A. & Lynch, S. V. Community ecology as a framework for human microbiome research. Nat Med 25, 884–889 (2019).

33. Prach, K. & Walker, L. R. Four opportunities for studies of ecological succession. Trends in Ecology & Evolution 26, 119–123 (2011).

34. Douglas, A. E. The microbial exometabolome: ecological resource and architect of microbial communities. Philosophical Transactions of the Royal Society B: Biological Sciences 375, 20190250 (2020).

35. Lee, J.-Y., Tsolis, R. M. & Bäumler, A. J. The microbiome and gut homeostasis. Science 377, eabp9960 (2022).

36. Van Herreweghen, F., De Paepe, K., Roume, H., Kerckhof, F.-M. & Van de Wiele, T. Mucin degradation niche as a driver of microbiome composition and Akkermansia muciniphila abundance in a dynamic gut model is donor independent. FEMS Microbiology Ecology 94, fiy186 (2018).

37. Ridlon, J. M., Kang, D. J., Hylemon, P. B. & Bajaj, J. S. Bile acids and the gut microbiome. Current Opinion in Gastroenterology 30, 332–338 (2014).

## METHODS REFERENCES

37. Howe, A. et al. Divergent responses of viral and bacterial communities in the gut microbiome to dietary disturbances in mice. ISME J 10, 1217–1227 (2016).

38. Carmody, R. N. et al. Diet dominates host genotype in shaping the murine gut microbiota. Cell Host and Microbe 17, 72–84 (2015).

39. Turnbaugh, P. J., Bäckhed, F., Fulton, L. & Gordon, J. I. Diet-Induced Obesity Is Linked to Marked but Reversible Alterations in the Mouse Distal Gut Microbiome. Cell Host and Microbe 3, 213–223 (2008).

40. Diaz-Ochoa, V. E. et al. Salmonella mitigates oxidative stress and thrives in the inflamed gut by evading calprotectin-mediated manganese sequestration. Cell Host Microbe 19, 814–825 (2016).

41. Bolger, A. M., Lohse, M. & Usadel, B. Trimmomatic: a flexible trimmer for Illumina sequence data. Bioinformatics 30, 2114–2120 (2014).

42. Eren, A. M., Vineis, J. H., Morrison, H. G. & Sogin, M. L. A Filtering Method to Generate High Quality Short Reads Using Illumina Paired-End Technology. PLOS ONE 8, e66643 (2013).

43. Li, D., Liu, C.-M., Luo, R., Sadakane, K. & Lam, T.-W. MEGAHIT: an ultra-fast single-node solution for large and complex metagenomics assembly via succinct de Bruijn graph. Bioinformatics 31, 1674–1676 (2015).

44. Eren, A. M. et al. Anvi’o: an advanced analysis and visualization platform for ’omics data. PeerJ 3, e1319 (2015).

45. Hyatt, D. et al. Prodigal: Prokaryotic gene recognition and translation initiation site identification. BMC Bioinformatics 11, 1–11 (2010).

46. Eddy, S. R. Accelerated Profile HMM Searches. PLOS Computational Biology 7, e1002195 (2011).

47. Bateman, A. et al. The Pfam protein families database. Nucleic Acids Research 32, D138–D141 (2004).

48. Kanehisa, M. & Goto, S. KEGG: Kyoto Encyclopedia of Genes and Genomes. Nucleic Acids Research 28, 27 (2000).

49. Zhang, H. et al. dbCAN2: a meta server for automated carbohydrate-active enzyme annotation. Nucleic Acids Res 46, W95–W101 (2018).

50. Langmead, B. & Salzberg, S. L. Fast gapped-read alignment with Bowtie 2. Nature Methods 2012 9:*4* 9, 357–359 (2012).

51. Li, H. et al. The Sequence Alignment/Map format and SAMtools. *Bioinformatics (Oxford*, England) 25, 2078–2079 (2009).

52. Wickham, H. et al. Welcome to the Tidyverse. Journal of Open Source Software 4, 1686 (2019).

53. Morrison, D. J. & Preston, T. Formation of short chain fatty acids by the gut microbiota and their impact on human metabolism. Gut Microbes 7, 189–200 (2016).

54. Devlin, J. R. et al. Salmonella enterica serovar Typhimurium chitinases modulate the intestinal glycome and promote small intestinal invasion. PLOS Pathogens 18, e1010167 (2022).

55. Heinken, A. et al. Genome-scale metabolic reconstruction of 7,302 human microorganisms for personalized medicine. Nat Biotechnol 41, 1320–1331 (2023).

56. Arkin, A. P. et al. KBase: The United States Department of Energy Systems Biology Knowledgebase. Nature Biotechnology 2018 36:*7* 36, 566–569 (2018).

57. Aziz, R. K. et al. The RAST Server: Rapid Annotations using Subsystems Technology. BMC Genomics 9, 75 (2008).

58. Henry, C. S. et al. Microbial Community Metabolic Modeling: A Community Data-Driven Network Reconstruction. J Cell Physiol 231, 2339–2345 (2016).

