## Supplemental Information Titles for "Diet outperforms microbial transplant to drive microbiome recovery post-antibiotics"

**Supplementary Discussion:** Intervention Day 28 results; body weight results from colonization resistance experiments.

**Supplementary Tables:**

**Table S1:** Analyses of microbial biomass by CFU plating.

**Table S2:** 16S analyses of microbiome recovery including comparisons of alpha and beta diversity across treatment groups and timepoints.

**Table S3:** Metagenomic analyses of functional diversity and redundancy across timepoints and RC-ABX and WD-ABX treatment groups.

**Table S4:** Differential abundance analysis of metagenomics data at the KO and pathway level.

**Table S5:** Comparison of metabolomics TMS panel and SCFA panel across treatment groups and timepoints. Information on internal standards used in metabolomic analysis.

**Table S6:** Analysis of select metagenomic gene abundances over time.

**Table S7:** Quantification of residual fecal antibiotic from Day 0 of recovery immediately after antibiotic cessation through Day 7.

**Table S8:** Metabolic modeling results including correlations between abundances of metabolites in a sample and the predicted capacity of the taxa in that sample to use them, as well as flux outputs from community-level flux-balance analysis for each treatment group and timepoint.

**Table S9:** Analyses of alpha and beta diversity across mice in all treatment groups at Day 14 of recovery in intervention experiments.

**Table S10:** Statistical comparisons of fecal and tissue *St* load, body weight, histopathology, and qPCR inflammatory gene expression across all treatment groups in colonization resistance experiments. Includes information on qPCR primers used for analysis.
