## Supplementary Discussion for "Diet outperforms microbial transplant to drive microbiome recovery post-antibiotics"

***Intervention experiment Day 28 results***

Although microbiome recovery was severely impaired for all treatment groups fed post-ABX WD_D_ at Day 14, by Day 28 these mice exhibited substantially improved ASV richness and an overall community composition that more closely resembled the untreated no-ABX controls on each diet (Figure S7). Similarly, the mice fed post-ABX RC_D_ that received WD_M_ microbes, which had exhibited only partial recovery of alpha diversity by Day 14, showed improved recovery of ASV richness by Day 28. These data suggest that while transplant of RC_M_ microbes immediately after cessation of antibiotics is sufficient to completely restore alpha diversity in mice on RC_D_, mice that either receive WD_D_ diet or WD_M_ microbes benefit from a second, later transplant.

These data carry the implication that in spite of our previously observed limited taxonomic and metabolomic recovery in mice on WD between Day 1 and Day 14 (Figures 1, 2), the gut environment in mice on WD changes from a state that is hostile to microbial transplant in the aftermath of antibiotics towards one that can accommodate engraftment and regrowth. As stated in the main text, this cannot be attributed to slower clearance of the antibiotics themselves in mice on WD (Figure S4). This environmental change may be facilitated by the regrowth of microbial biomass around Day 7, and the associated re-emergence of (limited) community complexity and syntrophy as predicted by our metabolic model. Perhaps even a limited increase in community complexity is sufficient to prevent domination by a single taxon that would otherwise exclude competitors, and therefore enables broader engraftment by transplanted taxa. Or perhaps these taxa promote metabolomic changes that were subtle or not measured in our targeted panel but which still play a significant role in community dynamics.

While these data do not affect our conclusion that an appropriate dietary resource environment is both necessary and sufficient for *rapid* and robust microbiome recovery, we stipulate that, as the resource environment may spontaneously change over time even without dietary intervention, such interventions may become less necessary depending on the character of those changes*.* Notably, such spontaneous changes are not sufficient to permit robust recovery on their own, given the sustained dysbiosis that we observed in our long-term experimental cohort (Figure S1). However, after two weeks, they appear sufficient to render the gut amenable to a second round of FMT. Taken together, we argue that dietary intervention remains the most fundamentally efficient and consistent way to promote endogenous microbiome recovery after antibiotic treatment, as well as to create a gut environment that is receptive to FMT.

***Body weight change in colonization resistance experiments***

We evaluated change in body weight after infection by *St* as a gross metric of host pathology (Figure S8D). Mice that were not infected did not exhibit any significant changes in body weight through t=96 hpi. Among infected treatment groups however, all experienced weight loss relative to their pre-infection baseline regardless of diet or antibiotic treatment status, and at t=96 hpi, were generally statistically indistinguishable from one another (Table S10).

We hypothesized that the substantial weight loss across all infected treatment groups, including those with significantly higher or lower *St* loads, may reflect systemic infection rather than localized lower GI infection. There are two established models of *St* infection: in the “typhoid model,” sufficient doses of *St* in mice that are not pre-treated with antibiotics establish systemic infection by invading the epithelial tissues of the ileum, disseminating to other tissues, and inducing a typhoid-like response^18^. In the “colonization resistance” model, even low doses of *St* in antibiotic pre-treated mice establish a persistent luminal lower-GI tract infection, inducing acute inflammation of the cecum and colon, diarrhea, weight loss, and severe host pathology^16^. Tissue dissemination is, in theory, specific to the typhoid model, whereas lower GI enteritis is specific to the colonization resistance model. Thus, we examined dissemination of *St* to the liver, mesenteric lymph nodes, and spleen, as well as levels in the ileum, cecum, and colon^55^. We observed detectable levels of *St* in all treatment groups for all GI regions and tissues assayed, confirming dissemination of *St* beyond the gut (Figure S8E). The colonization resistance phenotype in which the WD-ABX-INF group experienced greater infection loads than other treatment groups, however, was specific to the three GI regions (Table S10).

In combination with our histopathology and inflammatory gene expression assays, which showed that severe lower GI inflammation was limited primarily to the WD-ABX-INF and WD-PBS-INF treatment groups, these results suggest that although all infected treatment groups likely experienced some degree of systemic infection and consequent pathology and weight loss, the lower-GI infection and enteritis phenotypes were specific to the WD-ABX-INF treatment group. This supports the conclusion that prolonged post-antibiotic dysbiosis in mice on WD impairs colonization resistance in the lower GI tract relative to mice that were not pre-treated with antibiotics, or that were on RC diet.
